# Structure of the RAF1-HSP90-CDC37 complex reveals the basis of RAF1 regulation

**DOI:** 10.1101/2022.05.04.490607

**Authors:** Sara García-Alonso, Pablo Mesa, Laura de la Puente Ovejero, Gonzalo Aizpurua, Carmen G Lechuga, Eduardo Zarzuela, Clara M Santiveri, Manuel Sanclemente, Javier Muñoz, Mónica Musteanu, Ramón Campos-Olivas, Jorge Martínez-Torrecuadrada, Mariano Barbacid, Guillermo Montoya

## Abstract

RAF kinases are RAS-activated enzymes that initiate signalling through the MAPK cascade to control cellular proliferation, differentiation, and survival. Here, we describe the structure of the full-length RAF1 protein in complex with HSP90 and CDC37 obtained by cryo-electron microscopy. The reconstruction reveals a RAF1 kinase with an unfolded N-lobe separated from its C-lobe. The hydrophobic core of the N-lobe is trapped in the HSP90 dimer while CDC37 wraps around the chaperone and interacts with the N- and C-lobes of the kinase. The structure indicates how CDC37 can discriminate between the different members of the RAF family. Our structural analysis also reveals that the folded RAF1 assembles with 14-3-3 dimers, suggesting that after folding follows a similar activation as B-RAF. Finally, disruption of the interaction between CDC37 and the DFG segment of RAF1 unveils potential vulnerabilities to attempt the pharmacological degradation of RAF1 for therapeutic purposes.

## INTRODUCTION

The RAF family of kinases (A-RAF, B-RAF and RAF1, also known as C-RAF) are the direct KRAS effectors responsible for the activation of the MAPK signaling pathway(Matallanas et al., 2011). The three RAF proteins share a common modular structure consisting of three conserved regions (CR) (Figure 1a). CR1 encompasses a RAS binding domain (RBD), necessary for membrane recruitment, and a cysteine rich domain (CRD), CR2 contains important inhibitory phosphorylation sites participating in RAF activation, and CR3 contains the kinase domain (Figure S1), whose phosphorylation is crucial for activation(Matallanas et al., 2011). The physiological regulation of RAF kinases is intricate and involves several steps including dimerization, protein–protein interactions, phosphorylation and dephosphorylation events and conformational changes(Roskoski, 2010). In the absence of activating interactions with RAS, RAF proteins are thought to exist in a closed autoinhibited conformation, stabilized by a 14-3-3 dimer binding simultaneously to a phosphorylated N-terminal site and a C-terminal site. Interaction of RAF kinases with membrane bound RAS proteins bound to GTP leads to the activation of their kinase activity by a multistage process that requires phosphorylation at multiple sites within the N-terminal half of the protein, as well as dimerization of their kinase domains. The RAF kinases have restricted substrate specificity and catalyze the phosphorylation and activation of MEK1 and MEK2, their downstream effectors within the MAPK pathway(Roskoski, 2010). Finally, oncogenic activation of these proteins by missense mutations or by generation of fusion proteins have been described in multiple tumor types (Holderfield et al., 2014).

**Figure 1.**
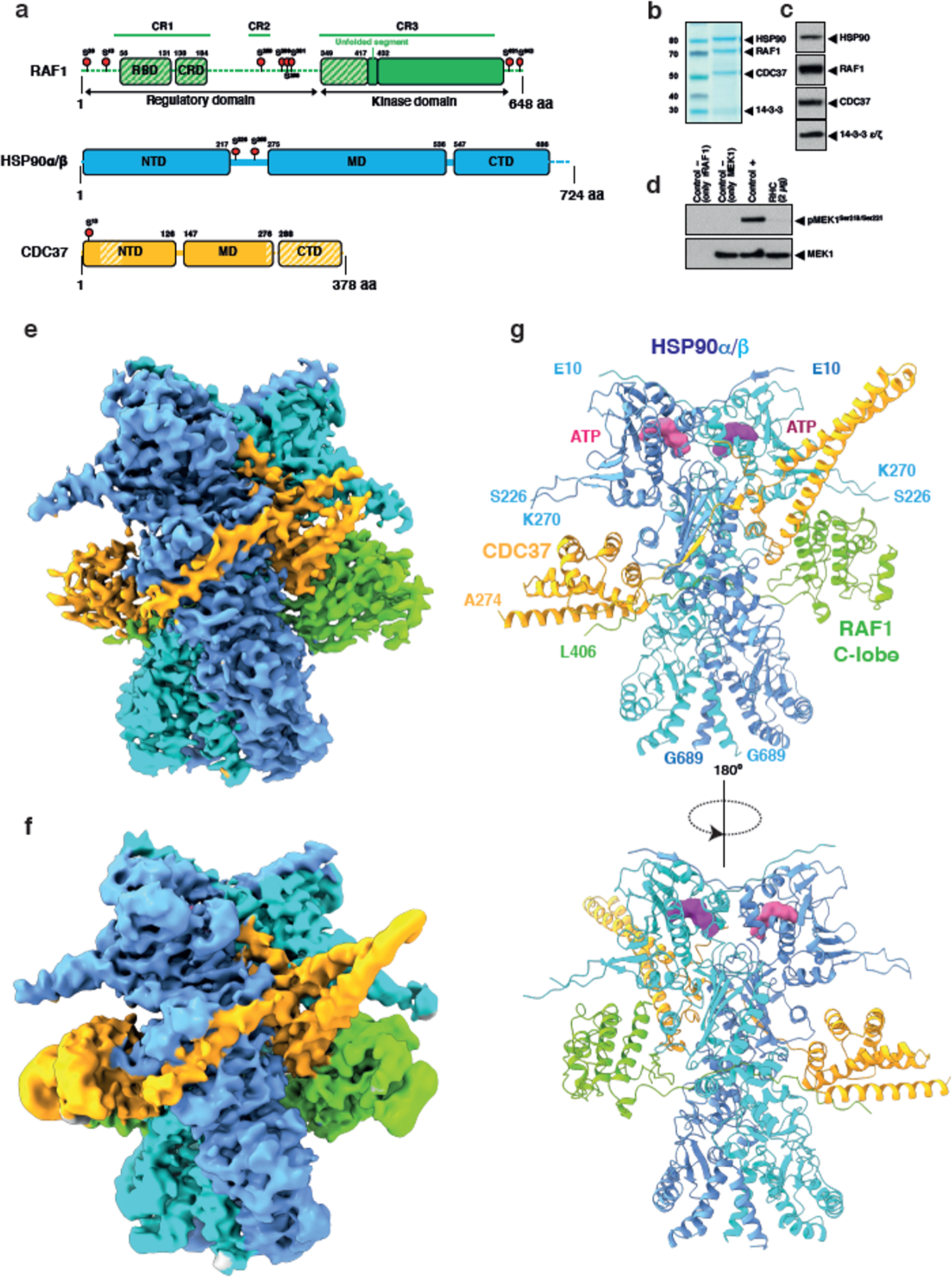
Biochemical characterization and cryo-EM structure of the RAF1-HSP90-CDC37 (RHC) complex. **a)** Schematic diagram showing the domain organization of the proteins present in the RHC complex. The stripped/dashed parts indicate regions where there is no interpretable density. Red circles indicate phosphorylated residues identified by mass spectrometry (See also Table S1). **b)** Coomassie-stained SDS-PAGE showing the affinity purified complex from Expi293F mammalian cells. Endogenous 14-3-3 proteins copurified with the complex (See also Figure S2). **c)** Western blot analysis to confirm the protein composition of the isolated fractions. Migration of the corresponding proteins is indicated by solid arrowheads. **d)** *In vitro* protein kinase assay using recombinant MEK1 protein as a substrate. **e-f)** cryo-EM maps of the RAF1-HSP90-CDC37 complex at 3.1 and 3.7 Å global resolution, respectively. The map is colored according to each subunit as in Figure 1a (See also Figures S3, S4, Table S2 and Movie S1), including blue and cyan for each monomer of the HSP90 dimer. **g)** Overview of the RAF1-HSP90-CDC37 assembly in complex with ATP (Surface representation colored in pink and magenta).

Based on their key role in MAPK signaling, RAF kinases have been long considered potential targets to block *KRAS* mutant tumors. Unfortunately, none of the different RAF inhibitors taken to the clinic, mainly kinase and dimerization blockers, have gone beyond phase II studies (Ryan and Corcoran, 2018). On the other hand, recent observations using genetically engineered mouse tumor models of *Kras/Trp53* driven lung adenocarcinoma have demonstrated that ablation of RAF1, but not of B-RAF or A-RAF, results in significant levels of tumor regression without inducing unacceptable toxicities (Sanclemente et al., 2018). Moreover, expression of kinase dead isoforms of RAF1 does not result in therapeutic consequences, indicating that the contribution of RAF1 to the progression of KRAS mutant lung adenocarcinomas is not mediated by its kinase activity (Sanclemente et al., 2021). These results strongly suggest that therapeutic strategies based on inhibition of RAF1 kinase activity are unlikely to produce anti-tumor results. Interestingly, RAF1 ablation does not affect MAPK signaling, suggesting that RAF1 contributes to tumor progression by other mechanisms, mainly inactivation of apoptotic pathways (Mikula et al., 2001; Sanclemente et al., 2021). For yet unknown reasons, RAF1 ablation does not induce tumor regression in mice harboring pancreatic tumors induced by the same *Kras/Trp53* mutations. Yet, combined ablation of RAF1 and EGFR results in the complete regression of a subset of these tumors. Hence, these results strongly suggest that therapeutic strategies based on inhibition of RAF1 kinase activity are unlikely to produce anti-tumor results (Blasco et al., 2019). Hence, novel therapeutic approaches that results in the pharmacological degradation of RAF1 should be considered.

RAF1 is found in both the cytosolic and membrane fraction of cells as a component of a large multisubunit protein complex, including the molecular chaperone HSP90 and the cochaperone CDC37 (Wartmann and Davis, 1994). The formation of the RAF1-HSP90-CDC37 (RHC) complex is crucial for RAF1 activity and MAPK pathway signaling (Grammatikakis et al., 1999), as the HSP90-CDC37 system facilitates RAF1 S621 phosphorylation and prevents protein degradation (Mitra et al., 2016). More than half of the human kinases are clients of the HSP90 molecular chaperone and its CDC37 cochaperone, who stabilize and activate them (Taipale et al., 2012). The structure at 4.0 Å of the CDK4-HSP90-CDC37 complex provided a first glimpse of the interaction of this chaperone system with a client kinase (Verba et al., 2016). However, this data does not provide the basis to understand why RAF1 and A-RAF, but not B-RAF, are part of its clients, and how this interaction affects RAF1 regulation.

Here, we provide evidence illustrating why the HSP90-CDC37 system can discriminate between its clients RAF1 and A-RAF but does not “chaperonise” wild type B-RAF. CDC37 recognizes specific regions in the C-lobe of RAF1 and mutations in the main interacting regions of CDC37 and RAF1 dissociate the RAF1-HSP90-CDC37 complex. Furthermore, cell lines including these mutations display a reduced RAF1 content and cell progression. Both can be restored after expression of the wild type RAF1, thus suggesting that disruption of this ternary complex represents a structural vulnerability that could render a therapeutic target for *Kras/Trp53*-driven tumors.

## RESULTS

### Isolation of the RHC complex

The RHC complex was isolated after transient co-expression in eukaryotic Expi293F cells (Figure 1b-c, Figure S2). This complex consists of four polypeptides including an HSP90 dimer, and one molecule of both CDC37 and RAF1. As determined by Western blot analysis and mass spectrometry, a sub-stoichiometric amount of 14-3-3 proteins mainly, but not exclusively, composed of the ε/ζ isoforms, copurified with the RHC complex (Figure S2d). The isolated assembly shows phosphorylations in RAF1 key residues as well as in HSP90 and CDC37 (Figure 1a, Table S1). Yet, we could not observe any density for the 14-3-3ε ζ that the key S259 and S621 residues of RAF1 are phosphorylated (Figure 1e-f, Figure S3-S4), suggesting that these dimeric phospho-binding proteins do not associate with RAF1 when it is coupled with the HSP90-CDC37 chaperone system. In addition, the isolated complex did not significantly phosphorylate MEK1 (Figure 1d), indicating that RAF1 in complex with the HSP90-CDC37 system is inactive.

### Overall structure of the RHC complex

Next, we aimed to obtain structural information of the RHC complex by cryo-EM to understand how the HSP90-CDC37 system recognizes RAF1. The purified complex was isolated in the presence of molybdate to stabilize the substrate for cryo-EM analysis (Hartson et al., 1999). Using a combination of statistical classification and Movie processing, we obtained two maps at 3.1 and 3.7 Å global resolution by the Fourier shell correlation (FSC)=0.143 criterion (Figure 1e-f, Figure S3-S4, Table S2). Both maps were combined during model building to generate an atomic model of the RHC complex. Computational sorting of images through regularized likelihood optimization and 3D variability analysis (Punjani and Fleet, 2021) allowed us to determine a variety of conformations present in the same sample (Scheres, 2012). The 3D variability analysis resolved the continuous flexibility of the sample representing different conformations of the assembly (Movie S1), thus illustrating the large plasticity of the RHC complex (Figure 1e-f, Figure S3-S4 and Table S2). The core of the RHC complex, specially HSP90, was visualized at high resolution; while the CR1 and CR2 regions of RAF1, as well as part of the N-terminal lobe of the kinase domain and the N- and C-terminal domains (NTD and CTD) of CDC37 were not visible in these density maps (Figure 1a, e-g), indicating the large mobility of these regions. In addition, changes in the conformation of the identified domains could be also observed between the different maps (Figure 3, Movies S2 and S3). The two maps displayed unambiguous densities for the protein side chains in the central section of the complex, while the peripheral regions display lower resolution and thereby less well-resolved 3D features, depending on their level of flexibility. The combination of both maps allowed us to generate a complete model (Figure 1e-g, Figure S3-S4). Nevertheless, the maps were of sufficient quality to accurately build and refine an atomic model of the RAF1-HSP90-CDC37 complex (Table S2), which permitted a detailed visualization of the interactions within the assembly.

### HSP90 separates the N- and C-lobes of RAF1

In the RHC complex structure, the HSP90 dimer adopts a conformation that closely resembles its closed state, as previously observed in the CDK4-HSP90-CDC37 structure (Verba et al., 2016). A segment of RAF1 is associated with a previously mapped HSP90 client binding site. The N-lobe of RAF1 is substantially unfolded, and it is visualized only in low resolution maps where it displays a large mobility (Figure 3c). Inside the core of HSP90, we observed an elongated polypeptide comprising part of the N-lobe and the connection with the C-lobe of RAF1 kinase domain (Figure 2a-b). The large range of flexibility of RAF1 in association with the HSP90-CDC37 chaperone system is shown in one of the classes, where only the section of RAF1 engaged with HSP90 could be observed (Figure 3a). This segment (L406 to W429) encompasses the complete β4 and β5 sheets, the connecting loops and the first turn of the αC helix of the RAF1 kinase domain. This unfolded region constitutes the core of the RAF1 kinase N-lobe associated with HSP90. Except the initial turn of the αD helix, the rest of the C-lobe of RAF1 is folded and can be superimposed with the corresponding region of the RAF1 crystal structure with only 1.2 Å RMSD (151 Cα atoms, PDB:3OMV) (Figure S5a). Collectively, the data indicates the N-lobe of RAF1 is largely flexible in our structure because it cannot form a folded domain when is associated to HSP90-CDC37.

**Figure 2.**
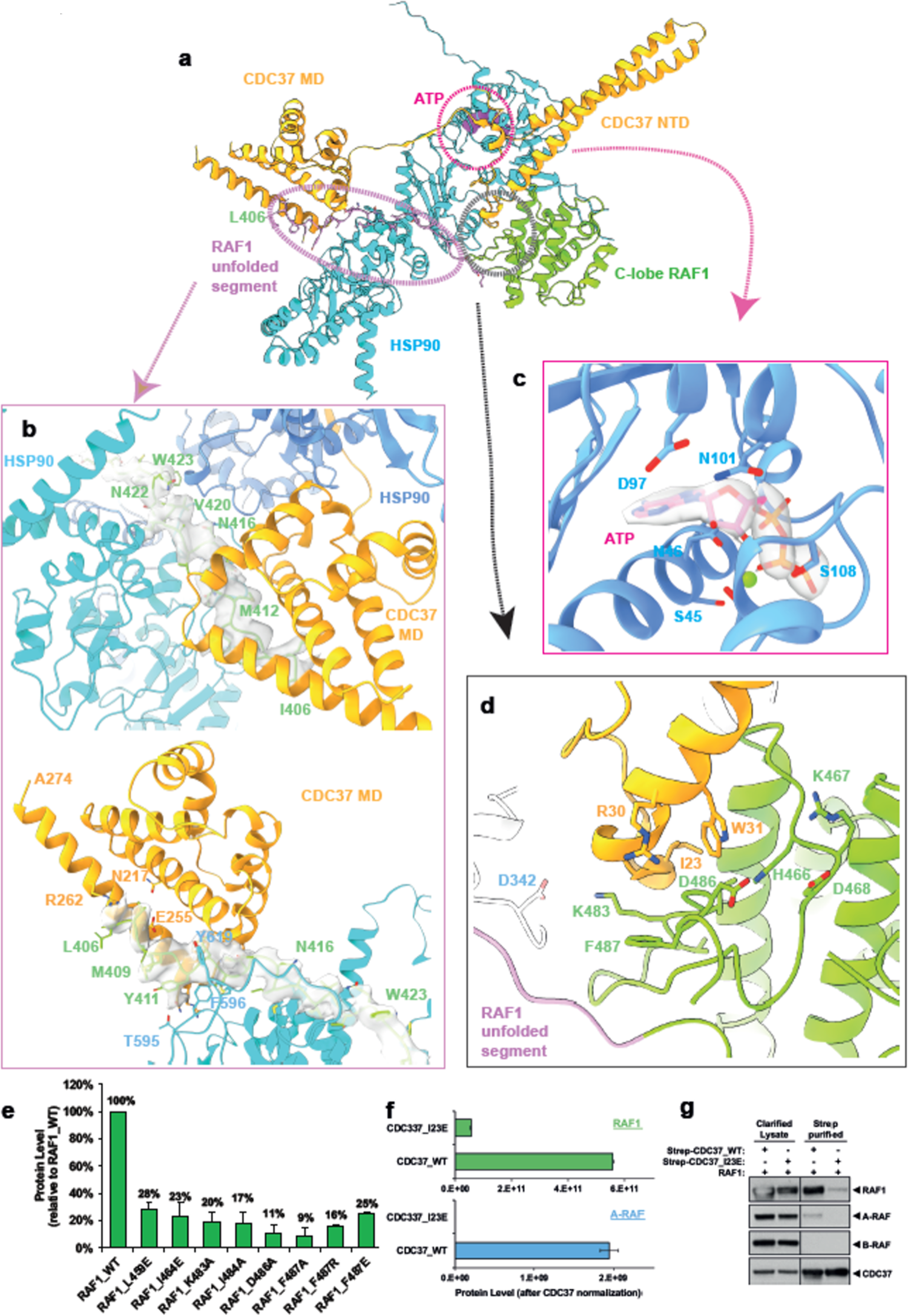
Interactions of RAF1 with HSP90 and CDC37 in the RHC complex. **a)** Overall view of the RHC complex. A monomer of HSP90 is omitted to show the region of RAF1 passing through the chaperone lumen and its interaction with CDC37. **b)** The top panel shows a view of the cryo-EM map of the L406-L429 region of RAF1. The bottom panel depicts a view of the L406-L417 region of RAF1 that interacts with the MD/CTD domain of CDC37 (See also Figure S5). **c)** View of the cryo-EM map around one of the ATP molecules at the NTD of HSP90. **d)** Detailed view of the main interacting region between CDC37 and RAF1 in the RHC complex. **e)** Levels of wild type and mutant RAF1 proteins co-expressed with HSP90 and CDC37 in Expi293F cells after strep-tag isolation from clarified cell lysates. The levels of each protein were quantified by mass spectrometry, using RAF1 common peptides for all the mutants. The graph shows the mean protein level, normalized to RAF1 wild type, of three different experiments. **f)** Mean levels of RAF1 and A-RAF bound to strep-tagged wild type CDC37 and mutant CDC37 I23E proteins. B-RAF peptides were undetectable (not shown). **g)** Western blotting analysis of clarified cell lysates and pull-downs from the experiment described in F. Note that the endogenous A-RAF protein co-purified with wild type but not with mutant CDC37. Migration of the corresponding proteins is indicated by solid arrowheads.

**Figure 3.**
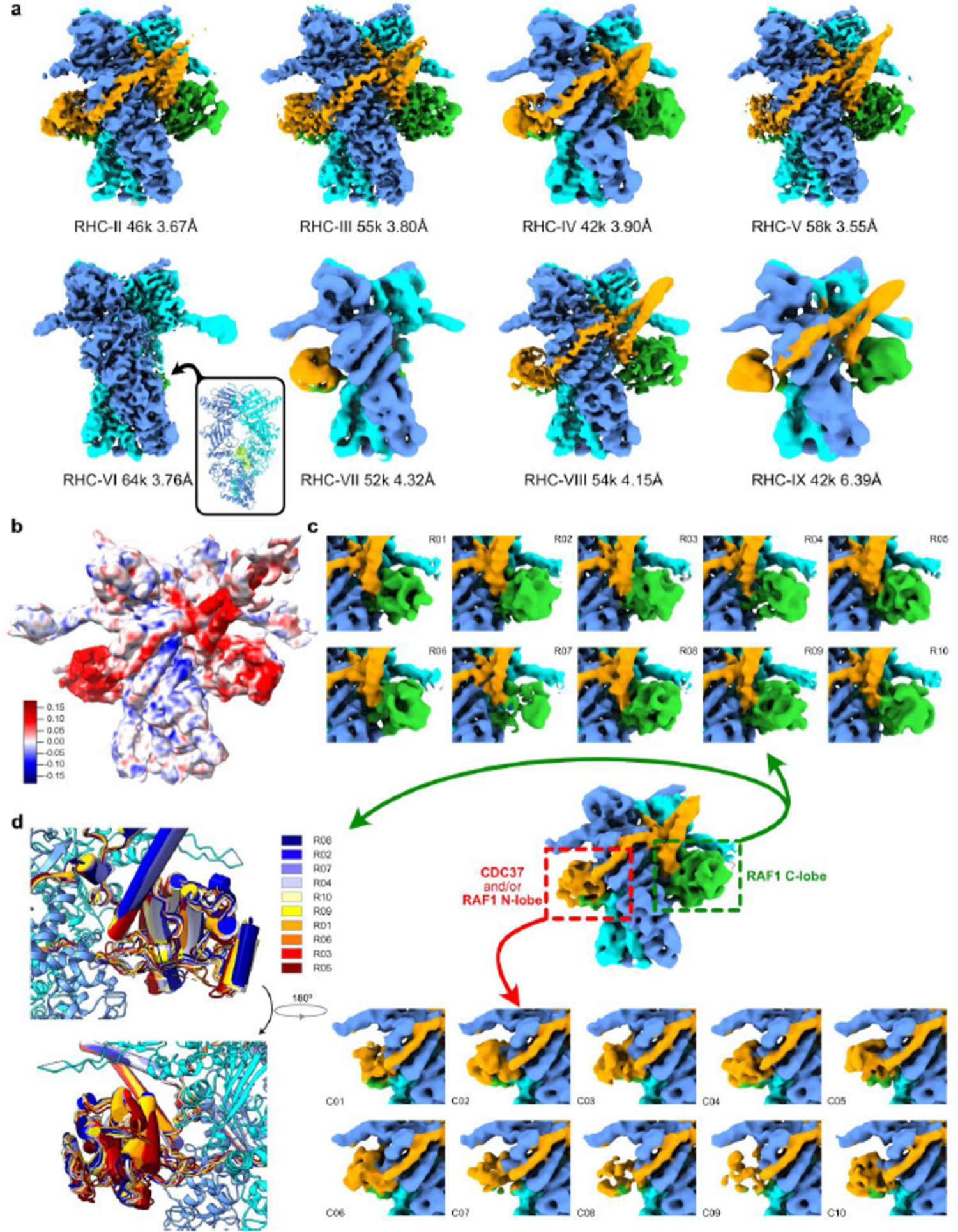
Conformational heterogeneity of the RHC complex. **a)** Eight different maps obtained from a single 3D classification, showing that while the HSP90 dimer constitute a stable core, the densities associated with RAF1 C-lobe and CDC37 (and/or RAF1 N-lobe) are significantly heterogeneous. All the maps and sections of densities or models in the whole Figure are displayed in the same orientation and colored according to the different subunits, as indicated in Fig 1. The number of particles and the global resolution of each map is indicated. The inset shows a lateral view of RHC-VI where only HSP90 (cartoon) and the density (in green) of the segment of RAF1 inside the lumen of HSP90 can be observed. **b)** 3D variability analysis of a subset of 442k particles of the RHC complex (cryoSPARC). The 3D reconstruction obtained from those particles is colored according to one dimension of the variability (from blue to red) (See also Movie S1). **c)** Local 3D classification of the previous subset of particles (See Figure S3c) focused on the RAF1 C-lobe region (**upper panel**) or the CDC37 MD and/or RAF1 N-lobe region (**lower panel**). Although these maps reach a global resolution range of 3.7 to 3.9 Å, they were filtered to 6 Å in this Figure for the sake of clarity, as the focused regions show lower local resolution values than the HSP90 core (See also Figure S4a and S4f). **d)** RAF1 C-lobe models were fitted and refined into the different locally focused maps, showing that the heterogeneity in that region can be described as the free rotation of the C-lobe around a fulcrum defined by its interaction with CDC37, and the entry point into HSP90 of the extended segment of RAF1 N-lobe (See also Movies S2 and S3).

The RAF1 residues ranging from W429 to L417 interact with the binding pocket of the HSP90 via extensive hydrophobic interactions (Genest et al., 2013). Only a polar interaction between R612, in one of the HSP90 chains, and RAF1 E425 at the entrance of the HSP90 cleft, can be observed (Figure 2a-b, Figure S5b). The HSP90 monomers collaborate in stabilizing the elongated hydrophobic segment; however, one of the HSP90 monomers displays a larger number of hydrophobic contacts, revealing an asymmetric interaction of the unfolded kinase lobe with the chaperone. This asymmetric association is related to the proposed mechanism used by the MD-CTD interface of HSP90 to link its nucleotide hydrolysis state to the associated client (Lavery et al., 2014).

The HSP90 nucleotide binding pockets are occupied in both maps, which were refined without 2-fold symmetry. Although molybdate is proposed to act as a post hydrolysis transition state inhibitor, thus helping to stabilize the HSP90 closed conformation, its mode of action is unclear (Hartson et al., 1999). Our results suggest that molybdate stabilisation could not be related to the nucleotide hydrolysis. The highest density value in the maps of the RHC complex is observed in the β- and γ-phosphates of the ATP molecule, suggesting that the complex traps HSP90 in an ATP bound state, as the bond between both phosphates appears to be intact (Figure 2c, Figure S3-S4). Therefore, the presence of molybdate during the purification may have stabilized the assembly independently of the locking of the post-hydrolysis transition state.

### CDC37 interacts with the N- and C-lobes of RAF1

The CDC37 NTD and middle domain (MTD)/CTD are also separated by the HSP90 dimer. The helical N-terminal contains a leucine zipper like motif and colocalizes with the C-lobe of the kinase. This region is linked with the CTD by a β-strand which flanks the HSP90 MD of one monomer, building an antiparallel β-sheet with the MD of this monomer. The β-strand is connected to the C-terminal density on the other side of the HSP90 dimer where CDC37 makes contacts with the residues 406-415 of RAF1 (Figure 1g, 2a-b, Figure S5c). Using a combination of the two maps, we were able to unambiguously resolve the structure until residue S300 of CDC37, thus visualizing the interaction of this region with the N-lobe of RAF1. Although distinct densities connected to the unfolded N-lobe are visible, and part of the N-lobe of RAF1 could be visualized, we did not model it, as we did not expect that this region of RAF1 could form the N-lobe due to the association of its core with HSP90 (Figure 2a, Figure 3a-c).

The C-lobe of RAF1 is stabilized by a combination of polar and hydrophobic contacts with the helical region of the NTD of CDC37. The residues T19 to R32 in CDC37 engage in interactions with the activation loop of RAF1, including the DFG and HRD motifs and the residues Q451 to I465 in the region of the αC and αD helices of RAF1 (Figure 2d, Figure S5d-e). The loop between residues T19 to T25 in CDC37 superimposes with the loop between αC and β4 of the RAF1 kinase domain structure, suggesting that the CDC37 NTD associates with the C-lobe by mimicking the interactions with a folded RAF1 N-lobe, thus stabilizing the separated C-lobe moiety. Our cryo-EM analysis permits the identification of these stabilizing interactions of the HSP90-CDC37 complex with the elongated RAF1 kinase domain. In addition, it shows that both, this association, and the globular C-lobe are flexible (Figure 3a-c, Movie S2, S3). Hence, indicating the mobility of certain domains of RAF1 when associated to the HSP90-CDC37 complex.

The phosphorylation of S13 in CDC37 is important to promote kinase function (Vaughan et al., 2008). We visualise the phosphate group in S13, which makes polar interactions with R36 and H33 in CDC37 and K406 in the corresponding monomer of HSP90 (Figure S5f). The phosphorylation stabilizes the N-terminus of the helical NTD of CDC37 and thereby favors the β-strand with HSP90. The arrangement promoted by S13 phosphorylation in the RHC complex is similar to that observed in the CDK4/HSP90/CDC37 complex (Verba et al., 2016), suggesting that this phosphorylation is a conserved mechanism of the HSP90-CDC37 system to promote the conformation leading to the interaction with the client independently of the kinase sequence (Verba et al., 2016).

After passing through the HSP90 lumen, the D415 to I406 section of the unfolded RAF1 N-lobe associates with the MD/CTD of CDC37 and engages in hydrophobic and polar interactions with residues M606, M620, Y596, Y619 and T595 with the HSP90 in one monomer. The D245, R246, Q247 residues in CDC37 are also involved in that interaction. The rest of the visualized unfolded N-lobe, ranging from Y411 to I406, lies on a cavity made by two helices where it engages in polar contacts with R262, E255, N217 and K257 residues in CDC37 (Figure 2a-b). The high mobility in this area hindered the observation of the rest of the N-lobe and the CTD of CDC37 (Figure 3).

### Recombinant 14-3-3τ protein promotes the dissociation of the RHC complex

As indicated above, RAF1 is phosphorylated in S259 and S621 in the RHC complex (Figure 1a, Table S1). These residues are equivalent to S365 and S729 in B-RAF, that are associated to 14-3-3 proteins in different B-RAF complexes(Kondo et al., 2019; Park et al., 2019). Although 14-ε/ζ proteins copurified with the RHC complex we could not detect stable complexes of RAF1 with these phosphate-binding proteins. To test whether 14-3-3 proteins could bind to the phosphorylated RAF1 present in RHC, we incubated recombinant 14-3-3ζ with the isolated RHC complex and analyzed the sample by cryo-EM. While a substantial fraction of the observed particles still belongs to the RHC complex, a new type of particles appeared in the sample and could be separated *in silico.* The reconstruction of the new particles at 7 Å allowed us to model a dimer of the RAF1 kinase associated to a 14-3-3 dimer using their crystal structures (fig S3, S6). This structure is similar to those of B-RAF/14-3-3 complexes (Kondo et al., 2019; Martinez Fiesco et al., 2022), which have provided an explanation for the mechanism of activation of B-RAF. Therefore, our observations suggest that the high concentration of 14-3-3 in the sample promotes binding of the phosphorylated RAF1 protein upon its release from the RHC complex. Whether RAF1 is folded by the chaperones prior to its release or after binding to 14-3-3, remains to be determined. However, the fact that RAF1 is stabilized by the HSP90-CDC37 strongly indicates that association to the phosphate-binding proteins occurs when the kinase domain is folded. Therefore, our data suggest that RAF1 activation may follow a similar mechanism as B-RAF, after its folding by the HSP90-CDC37 system. Unfortunately, the low resolution of the map does not allow us to visualize if the RAF1 dimer is arranged in a way that one kinase monomer can activate the other, as proposed for B-RAF (Kondo et al., 2019).

### Disruption of the interaction between CDC37 and RAF1 affects protein stability

To interrogate the functional relevance of the interactions observed within the structure of the RHC complex, we isolated complexes formed between eight different RAF1 mutants co-expressed with HSP90 and CDC37 in Expi293F cells. The relative levels of the mutant RAF1 proteins in the isolated RHC complexes were assessed by mass spectrometry and compared to those formed by the wild type protein. Mutation of key residues in the interface between RAF1 and CDC37 reduced the level of RAF1 protein present in the complexes between 70 to 90% (Figure 2e). Likewise, expression of a CDC37 protein carrying the I23E mutation, a residue located in the loop that stabilizes the C-lobe of RAF1 (Figure 2d), with wild type RAF1, strongly affected the amount of RAF1 present in the RHC complex. Similar results were observed for the endogenous A-RAF protein, highlighting the importance of this residue in the stabilization of the C-lobe (Figure 2f-g, Methods). As expected, B-RAF did not copurify with CDC37 in any case. Collectively, the site-directed mutagenesis data supports the importance of the interactions observed between CDC37 and the C-lobe of RAF1 and suggest that the client kinase is degraded when its association with the HSP90-CDC37 system is disturbed.

### Structural basis for the HSP90-CDC37 discrimination between RAF proteins

The interactions observed within the RHC complex prompted us to interrogate the molecular bases for the different association of the three RAF proteins with the HSP90-CDC37 system. A previous study concluded that the binding determinants of the RAF kinases to HSP90 are widely distributed in both lobes of their kinase domain and act in a combinatorial way to determine the interaction (Taipale et al., 2012). Our structure shows that the binding of HSP90 stabilizes the hydrophobic core of RAF1 N-lobe, as previously shown for the CDK4/HSP90/CDC37 complex(Verba et al., 2016), while CDC37 seems to recognize different regions of the RAF1 kinase domain and combines the non-polar contacts with polar interactions.

To test whether CDC37 can recognize specific segments of the various RAF kinases, we performed a binding assay incubating this cochaperone with a nitrocellulose-bound dodecapeptide array (PepScan) displaying the corresponding sequences of RAF1, B-RAF and A-RAF kinase domains (Figure 4). The assay shows that CDC37 can bind and is able to discriminate between the different dodecapeptides in the absence of HSP90. Noteworthy, CDC37 does not bind the region of RAF1 interacting with HSP90. Instead, the array shows that, despite some sequence differences, CDC37 recognizes a region in the N-lobe common to the three RAF family members (residues M350-W368, RAF1 numbering) including the GSGSFG motif (Terasawa et al., 2006). In addition, CDC37 interacts preferentially with another segment of RAF1 (P384-T400), not conserved in the other RAF proteins. This segment includes the loop joining αC with β4 (A395-T400) (Prince and Matts, 2004), which is found stabilizing the C-lobe of the unfolded RAF1 kinase domain (Figure 2a, d, Figure S5d-e), as previously observed in CDK4(Verba et al., 2016). Residues V482 to G498 display another high intensity PepScan binding site common to the three members of the RAF family. Finally, the C-lobe of RAF1 shows two specific binding sites in the P524-I536 and M560-K572 segments, which display minor sequence differences (Figure 4b-c, Figure S1). Therefore, CDC37 binds to different regions within the members of the RAF family. While several of these segments are shared among the three RAF proteins, the CDC37 cochaperone also associates with specific regions present in the RAF1 kinase domain, suggesting that the combination of the common and the exclusive sites is used by CDC37 to discriminate between the three family members.

**Figure 4.**
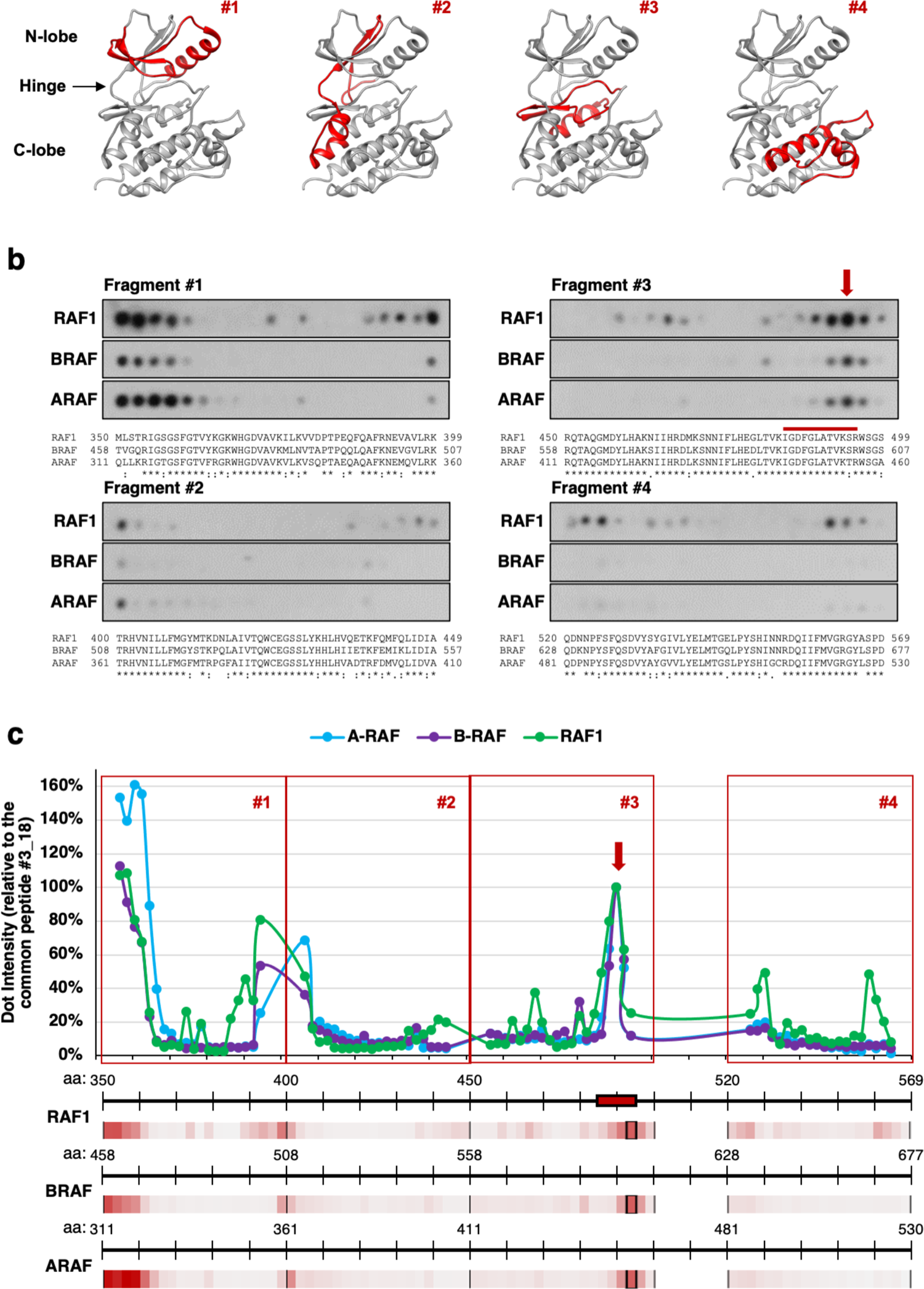
Recognition of RAF sequences by CDC37. **a)** Schematic diagrams showing the general structure of the kinase domain of RAF family members. The four fragments analysed by PepScan, covering the whole kinase domain, are highlighted in red (See also Table S3). **b)** CDC37 binding assay on nitrocellulose-bound RAF peptides (PepScan). Overlapping dodecapeptides with a shift of two amino acids residues scanning the indicated regions of the kinase domains of human RAF1, B-RAF and A-RAF proteins developed by ECL after incubation with purified CDC37. The sequences covered by these dodecapeptides are indicated below each set of membranes. **c)** Quantification of the PepScan assay for the sequences of the (green) RAF1, (purple) B-RAF and (blue) A-RAF kinase domains. The RAF1 peptide that interacts with CDC37 according to the cryo-EM structure is indicated with a red arrow. The CDC37 binding intensity to each peptide was calculated as the mean pixel integrated density of each dot relative to a reference one, with a common sequence among RAF family members (peptide 18 from region #3). The dot intensity is also shown as a colour-scale (light grey to red), with the reference dot marked by a black square.

### Disruption of the RAF1-CDC37 interface decreases RAF1 dependent proliferation

As illustrated above, the assembly of the RHC complex is severely reduced in D486A and F487A RAF1 mutants (Figure 2e). To evaluate the effect of these single mutations as well as the D486A/F487A double substitution in cell proliferation, we took advantage of *H-Ras^-/-^, N-Ras^-/-^, K-Ras^lox/lox^* (K-Ras^lox^) mouse embryonic fibroblasts (MEFs), which are known to cease proliferation upon ablation of KRAS expression (*Rasless* cells) ^26,27^ (Figure 5a). Ectopic expression of RAF1 in these cells restores cell proliferation through its interaction with the plasma membrane via addition of CAAX sequences derived from RAS proteins as long as p16INK4A expression is downregulated to avoid cell senescence (Drosten et al., 2010; Lechuga et al., 2021). We observed that the ectopic expression of wild type RAF1, but not the D486A, F487A and D486A/F487A mutants, elicited efficient proliferation of *Rasless* cells as determined in a colony-forming assay (Figure 5b-c). To validate this effect in cell proliferation, we use an additional cell model: *A-Raf^lox/lox^, B-Raf^lox/lox^, Raf1^lox/lox^* (Raf^lox^) MEFs, which also cease proliferation upon ablation of all RAF family members’ expression (*Rafless* cells) (Figure 5d). Ectopic expression of wild type RAF1 in these cells completely restored cell proliferation, albeit the absence of A-Raf and B-Raf. Contrarily, we observed that the ectopic expression of the mutant forms of RAF1 (D486A, F487A and D486A/F487A) did not promote proliferation of *Rafless* cells as determined in a colony-forming assay (Figure 5e-f). Thus, disruption of the interaction between the RAF1 mutants and CDC37 hinders cell proliferation most likely due to the instability of the RAF1 mutants.

**Figure 5.**
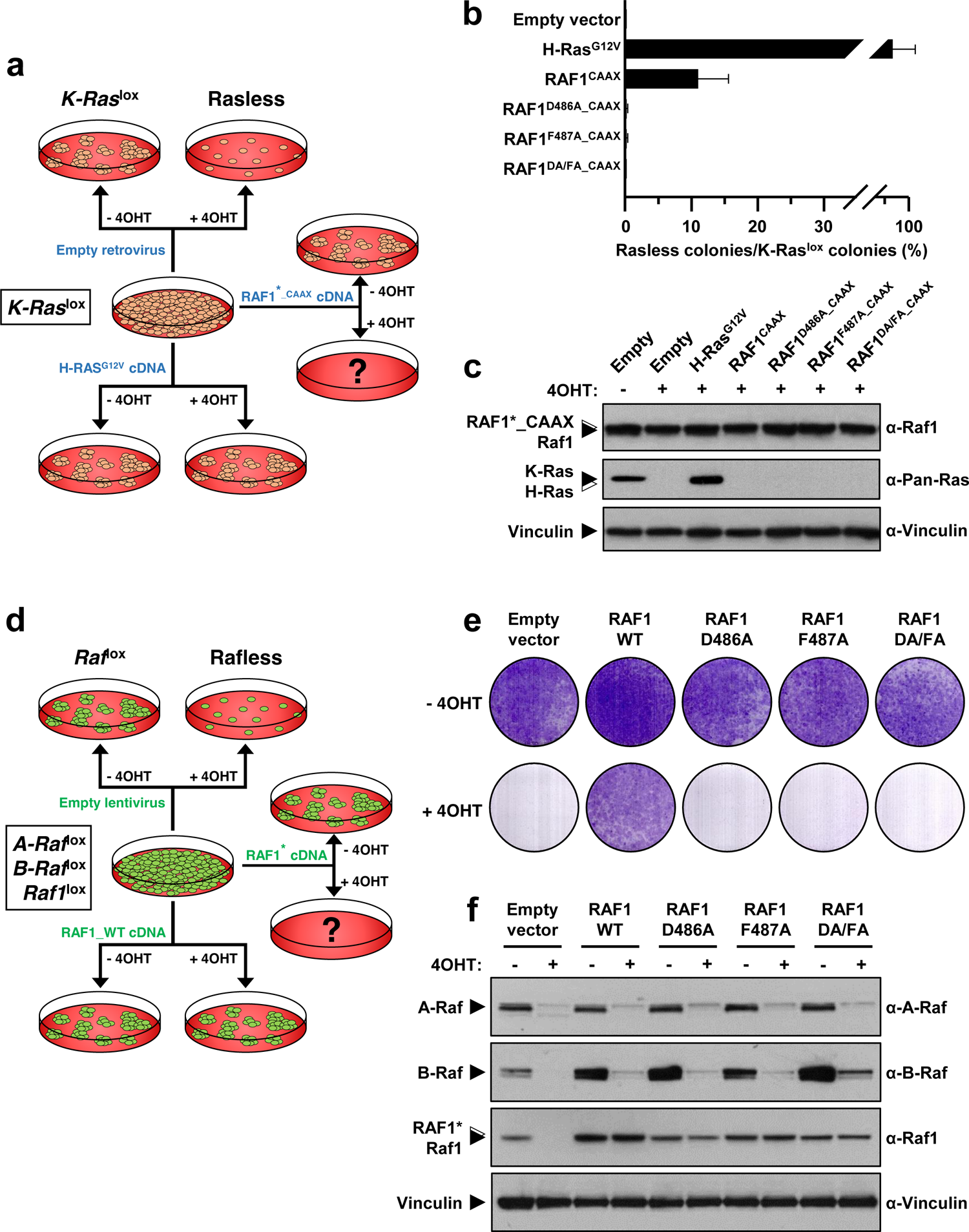
Disruption of the interaction between CDC37 and RAF1 inhibits cell proliferation. **a)** Schematic representation of the *Rasless* colony assay used to evaluate the capacity of RAF1 mutants to induce cell proliferation. **b)** Colony-formation assay using K-Ras^lox^ MEFs infected with retroviruses expressing the indicated cDNAs. Error bars indicate standard deviation. **c)** Western blot analysis showing levels of expression of the indicated H-Ras^G12V^ or RAF1 CAAX proteins in K-Ras^lox^ MEFs. Cells were either (**–**) left untreated or (**+**) exposed to 4OHT, to eliminate *Kras* expression. Probing antibodies are indicated in the right panel. Migration of the corresponding proteins is indicated by solid arrowheads. Open arrowheads indicate migration of the exogenous proteins. Vinculin expression served as loading control. **d)** Schematic representation of the *Rafless* colony assay used to evaluate the capacity of RAF1 mutants to induce cell proliferation. **e)** Colony-formation assay using Raf^lox^ MEFs infected with lentiviruses expressing the indicated cDNAs. **f)** Western blot analysis showing levels of expression of the indicated proteins in Raf^lox^ MEFs. Cells were either (**–**) left untreated or (**+**) exposed to 4OHT, to eliminate *A-Raf*, *B-Raf* and *Raf1* expression. Probing antibodies are indicated in the right panel. Migration of the corresponding proteins is indicated by solid arrowheads. Open arrowheads indicate migration of the exogenous proteins. Vinculin expression served as loading control.

Next, we interrogated whether disruption of the interaction between CDC37 and RAF1 had similar consequences in human tumor cells. To this end, we conjugated the top binding peptide (Figure 4b-c) to a TAT cell-penetrating motif (Green and Loewenstein, 1988) and added it to the A549 human lung cancer cell line and to cells derived from the Pdc1 lung cancer patient-derived xenograft (PDX), two cell lines known to be dependent on RAF1 expression for proliferation (Lee et al., 2013; Qiu et al., 2012; Sanclemente et al., 2018). As negative controls, we used a non-interacting RAF1 peptide and the HA peptide conjugated to TAT (Figure S7). Culture viability was measured by an MTT metabolization assay. The RAF1 interacting peptide selectively inhibited growth of both human lung cancer cells (Figure S7b). These observations suggest that the TAT derived RAF1 peptide may interfere with the RHC complex assembly, hindering RAF1 activity and thereby affecting cell proliferation.

## DISCUSSION

In this study, we visualize how RAF1 assembly with HSP90 and CDC37 is crucial for its stability and consequent activation(Grammatikakis et al., 1999) and prevents its degradation(Mitra et al., 2016), since the RHC complex maintains RAF1 in an inactive form. Thus, the open partially folded intermediate of RAF1 observed in the RHC complex plays a key regulatory role in controlling signaling through the MAP kinase cascade and cellular proliferation. Our data reveal that the structural principles of RAF1 kinase recognition by the HSP90-CDC37 system are similar to those observed in its complex with CDK4 (Verba et al., 2016). The complex displays a high flexibility, which is most likely indispensable when its function is to keep RAF1 kinase domain in a stable but quasi folded state (Figure 3). The mimicking of the loop joining αC and β4(Prince and Matts, 2004) in the kinase domain by CDC37 is a conserved feature of the HSP90-CDC37 complex, which explains the importance of the phosphorylation in the S13 residue of CDC37, as the phosphate group builds the interactions that promote the conformation to stabilize the folded C-lobe of RAF1. We also observed that a section of the unfolded N-lobe which emerges from the HSP90 binding site is stabilized by the MD/CTD region of CDC37.

Despite their well conserved sequence, members of the RAF family contain substantial functional differences. RAF1 and A-RAF are client proteins of the HSP90-CDC37 chaperone system, while B-RAF is not. Therefore, the HSP90-CDC37 chaperone system adds an extra regulatory layer to the members of this family. A key question in the intricate regulation of the RAF family is how the HSP90-CDC37 system can discriminate between them. Strong HSP90-CDC37 clients have been characterized to be less thermodynamically stable on their own, while more stable clients have reduced dependence on the chaperone system for maturation (Taipale et al., 2012). The structure of the RHC complex highlights the key interactions of the HSP90 chaperone and its cochaperone CDC37 with RAF1. Moreover, our combined biochemical and functional analysis of the interacting regions indicates that CDC37 can recognize segments of RAF1 that are different from their counterparts in B-RAF. In addition, the comparison of the crystal structures of RAF1 and B-RAF kinase domains suggest that the 480-505 region of RAF1, which is important in the interaction with CDC37, displays different structural features. This region is disordered in RAF1, while it is structured in B-RAF (Figure S8), suggesting that the sequence differences promote a different stability of this section of the kinase domain that interacts with CDC37. Therefore, our data suggest that discrimination between B-RAF and RAF1 by HSP90-CDC37 arise from the combined effect of the recognition of some specific sites of RAF1 by CDC37 and the different stability of the two kinase domains, which could make the main binding site preferentially available for CDC37 binding in the case of RAF1. Likewise, this may be the case for A-RAF, however, its kinase domain structure is not available for comparison.

This conclusion is also supported by the fact that the B-RAF V600E oncogenic mutant is a client of the HSP90-CDC37 system (Diedrich et al., 2017; Grbovic et al., 2006), which is in agreement with the proposed destabilization of the kinase inactive state by the V600E mutation by disrupting hydrophobic interactions present in the wild type, while the active state would be stabilized through the formation of a salt bridge between E600 and K507 residues (Maloney et al., 2021). This notion is also supported by the fact that the crystal structures of the B-RAF V600E mutant (PDB:4MNF, PDB:6P3D) display a disordered 480-505 region similar to that observed in RAF1 (Figure S8). Therefore, in the case of the B-RAF V600E, the HSP90-CDC37 system could assist the folding of the oncogenic mutant by promoting an active stable kinase which triggers malignant transformation. Collectively, our data suggest that recognition of the client kinase by CDC37 would depend on the binding sites of the target polypeptide and the availability of these sequences to associate with CDC37, which is controlled by the global stability of the protein.

The site directed mutagenesis analysis of the interface between CDC37 and the main binding region of RAF1 highlights the importance of this association for RAF1 stability. Indeed, we observed a strong reduction in the levels of RAF1 when the mutant isoforms were co-expressed with HSP90 and CDC37. In addition, germ line expression of a RAF1 D486A mutant in mice led to embryonic dead during mid-gestation due to the absence of RAF1 expression, a result likely to be due to the instability of this mutant protein (Noble et al., 2008). As illustrated here, RAF1 mutations in this interphase affect cell proliferation, since the mutant RAF1 proteins were not able to induce proliferation of MEF when ectopically attached to the plasma membrane in the absence of RAS proteins. Similar results were obtained using two independent human lung carcinoma cell lines treated with a peptide corresponding to RAF1/CDC37 interphase sequences, suggesting that this peptide interferes with the folding and hence, the activity of RAF1. These observations raise the possibility that the interphase between RAF1 and CDC37 may represent a vulnerable region, which could be targeted by small molecules to induce the degradation of RAF1. This potential pharmacological approach could reproduce the therapeutic results obtained in experimental models of *Kras/Trp53-*induced lung tumors upon ablation of RAF1 expression (Sanclemente et al., 2018).

We propose a model in which RAF1 would be unstable until it becomes associated with CDC37 followed by binding to HSP90, leading to the formation of the RHC complex (Figure 6). The HSP90-CDC37 chaperone system couples the folding of the client protein with ATP hydrolysis cycles (Figure 6a). RAF1 is phosphorylated in residues S259 and S621, thereby, once the HSP90-CDC37 renders this protein folded, the RHC complex is disrupted and RAF1 associates with 14-3-3 (Figure 6a). Although the low resolution cryo-EM structure of RAF1 associated with 14-3-3 limits a detailed interpretation, our data shows that the RAF1 protein associates with the dimeric adaptor proteins in a folded conformation of the kinase domain. The similarity of this structure with the B-RAF/14-3-3 complex (Kondo et al., 2019), and the phosphorylation sites found in the isolated RHC complex, suggest that the assembly of RAF1 with the HSP90-CDC37 system may facilitate the activating phosphorylation. Although the kinase domain will remain unfolded and therefore inactive, but this conformation could facilitate its phosphorylation in the S259 and S621 residues. Consequently, the RHC complex will maintain RAF1 in a “ready-to-act pool”. Once the folding of the kinase domain is completed, the 14-3-3 proteins would associate with the phosphorylated RAF1 to regulate its further activation (Figure 6a-b). On a different level of regulation, heterodimerization between RAF1 and B-RAF has been shown to be a part of the physiological activation process, and the heterodimer displays distinct biochemical properties likely to be important for signaling regulation (Rushworth et al., 2006). We speculate that the interaction of RAF1 with the HSP90-CDC37 system could control the dynamics of the RAF1/B-RAF heterodimers formed with the 14-3-3 proteins by providing folded and phosphorylated kinase, thus influencing the levels of homo or heterodimers of this signaling module, and thereby influencing cellular proliferation (Figure 6b).

**Figure 6.**
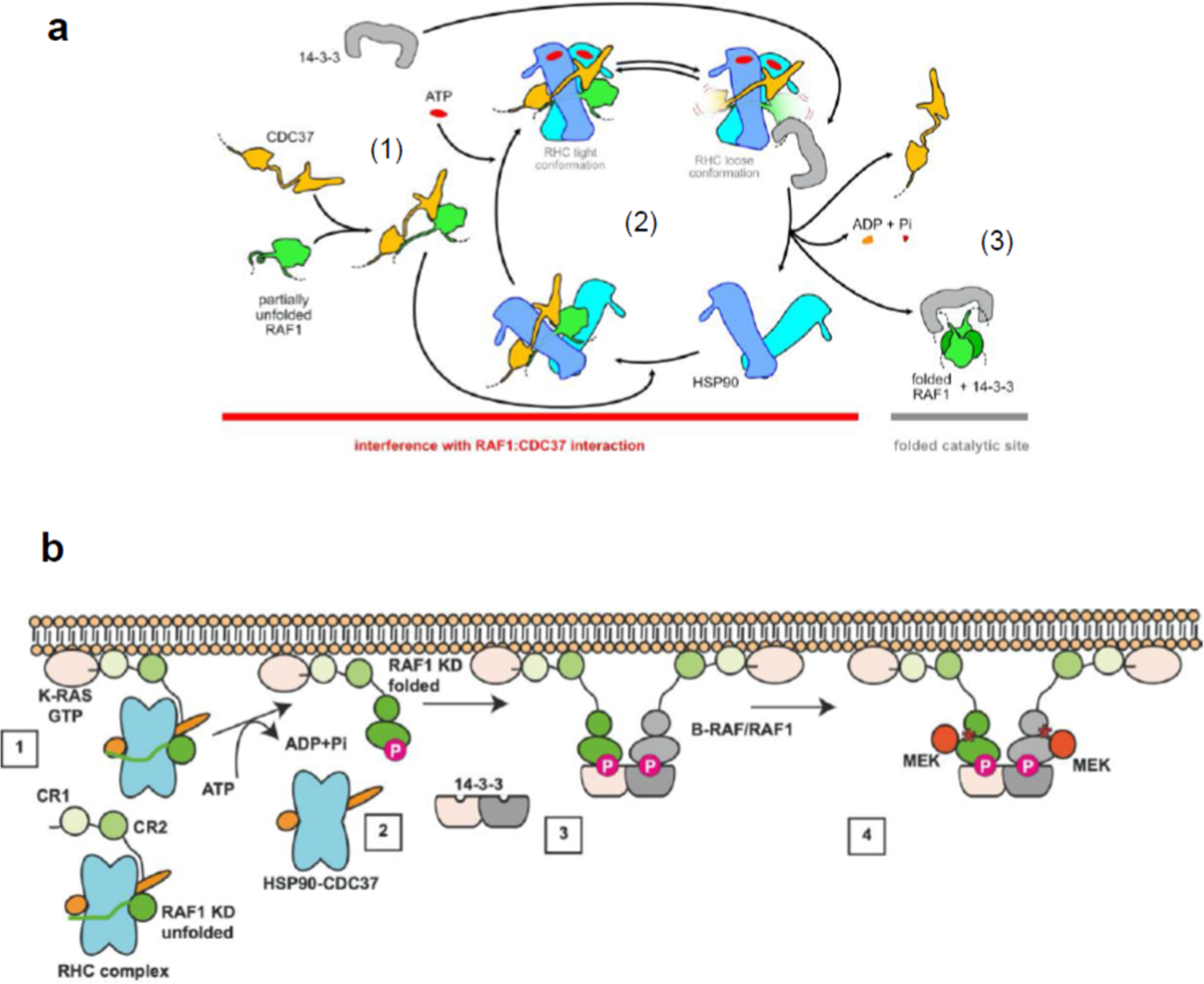
Model of RAF1 folding by the HSP90-CDC37 chaperone system and RHC function within the RAF1 regulation. **a)** The unfolded RAF1 protein associates to CDC37 and is stabilized after assembly with HSP90 forming the RHC (1). The RHC displays a large conformational heterogeneity to stabilize the unfolded kinase and proceeds to fold RAF1 coupling the conformational changes with the hydrolysis of ATP (2). The folded RAF1 can interact with 14-3-3 proteins through S259 and S621 (3). **b)** RAF1 is stabilized by the HSP90-CDC37 complex (1). After interaction with K-RAS in the GTP bound state and the hydrolysis of ATP by HSP90, the kinase domain of RAF1 is folded and S259 and S621 are phosphorylated (2). The folded and phosphorylated kinase binds 14-3-3 proteins, which could have the second binding site occupied with another RAF kinase, either RAF1 or B-RAF (3). The hetero or homodimer is activated (asterisk) modulating the activity and the phosphorylation of MEK depending on the assembly of RAF homo or heterodimers (4). Therefore, the downstream signaling will be controlled by the ratio of folded RAF1 available, which determines the proportion between homo and heterodimers, as B-RAF activity is not assisted by the HSP90-CDC37 system.

Given the contribution of RAF1 to *Kras/Trp53*-driven tumors, our work may open new avenues to address therapeutic approaches by either inhibiting the RHC complex formation or the recognition of RAF1 by CDC37. Yet, such therapeutic strategies would need to conserve the HSP90-CDC37 chaperone homeostatic function to avoid undesired toxicity.

## ACKNOWLEDGEMENTS

We thank the Danish Cryo-EM National Facility in CFIM at the University of Copenhagen for support during cryo-EM data collection. We also thank the Protein Expression Unit at CPR for assistance in protein expression and purification. S.G-A. is a recipient of a postdoctoral fellowship from the Spanish Association against Cancer Scientific Foundation (FCAECC). This work was supported by grants from the European Research Council (ERC-2015-AdG/695566, THERACAN); the Spanish Ministry of Science, Innovation and Universities (RTC-2017-6576-1), the Autonomous Community of Madrid (B2017/BMD-3884 iLUNG-CM), the CRIS Cancer Foundation and the AECC (GC16173694BARB) to M.B. and by a grant from the Spanish Ministry of Science, Innovation and Universities (RTI2018-094664-B-I00), to M.B. and M.M. M.B. is a recipient of an Endowed Chair from the AXA Research Fund. G.M. is part of the Novo Nordisk Foundation Center for Protein Research (CPR), which is supported financially by the Novo Nordisk Foundation (grant NNF14CC0001). This work was also supported by grant NNF0024386, grant NNF17SA0030214 and Distinguished Investigator (NNF18OC0055061) grants to G.M. G.M. is a member of the Integrative Structural Biology Cluster (ISBUC) at the University of Copenhagen.

## AUTHOR CONTRIBUTIONS

MB and GM conceived the study, SGA MB and GM designed the biochemical experiments. SGA and JM-T set up the purification protocol. SGA, LPO and GA created the mutants and performed biochemistry experiments. SGA and CGL performed tissue culture experiments, EZ and JM performed mass spectrometry analysis and analyzed the results, CMS and RC-O performed the SEC-MALS analysis, PM prepared EM grids and collected the cryo-EM images. PM performed cryo-EM data processing, built the models structure and PM and GM proceeded with cryo-EM map and structure analysis. The global results were discussed and evaluated with all authors. GM and MB coordinated and supervised the project and wrote the manuscript with input from all the authors.

## STAR METHODS

### Plasmid preparation and expression in Expi293F mammalian cells

The human full-length RAF1 cDNA (accession number NM_002880.3) was amplified by PCR with restriction site-tailed primers incorporating a C-terminal StrepTagII, separated from the protein by a Gly-Ser-Ala residue linker. The resulting DNA was cloned into the HindIII/XhoI sites of the transfer vector pcDNA3 (Invitrogen), using the In-Fusion® HD EcoDry Cloning Kit (Takara Bio USA). The derivative plasmid, pcDNA3-RAF1-Strep, was sequence verified. The plasmid encoding the full-length human HSP90 beta with a HA-tag at its N terminus was obtained from Addgene (Plasmid #22487). The cDNA clone for full-length human CDC37 tagged with a C-terminal Myc-DDK tag was purchased from Origene (RC201002). Expi293F cells (ThermoFisher Scientific) were maintained and expanded following the recommendations of the manufacturer. In short, Expi293F cells were grown in Expi293 Expression Medium (ThermoFisher Scientific) at 37 °C, 120 rpm, 5% CO_2_ atmosphere in PETG Erlenmeyer flasks with 0.2 μm ventilated caps (Corning) and split roughly every fourth day. Transfection of cells were performed with ExpiFectamine 293 Transfection Kit, following the manufacturer’s recommendations. For each transfection, cells were diluted to a final density of 3 mvc/ml in a total volume of 120 ml (in a 500 ml flask) and transfected with a total of 120 µg of plasmid DNA (56 µg of pcDNA3-RAF1-Strep + 32 µg of pcDNA3-HSP90-HA + 32 µg of pCMV6-CDC37-Myc-DDK), diluted with OptiMEM I-GlutaMAX (ThermoFisher Scientific). Cells were allowed to growth in standard conditions for 48 additional hours and then were harvested.

### Purification of the RAF1-HSP90-CDC37 complex

Four batches of transfected Expi293F cells (480 ml total) were resuspended individually in 15 ml of lysis buffer (20 mM Tris-HCl [pH 7.5], 150 mM NaCl, 10 mM MgCl_2_, 10 mM KCl, 20 mM Na_2_MoO_4_ and 0.1% Triton X-100) supplemented with complete Mini Protease Inhibitor Cocktail (Merck) and Phosphatase Inhibitor Cocktails 2 and 3 (Merk) and left on agitation at 4 °C for 15 minutes. After a brief sonication, the lysate was centrifuged (10,000 rpm, 10 min, 4 °C) and the soluble fraction was purified by affinity chromatography using a 5-ml StrepTrap® column (GE Healthcare), on an ÄKTA prime (GE Healthcare). The StrepTrap column was washed with 20 column volumes (CV) of buffer A1 (20 mM Tris-HCl [pH 7.5], 150 mM NaCl, 10 mM MgCl_2_, 10 mM KCl, 20 mM Na_2_MoO_4_) and eluted with a stepwise gradient (0–100%) of buffer B1 (20 mM Tris-HCl [pH 7.5], 150 mM NaCl, 10 mM MgCl2, 10 mM KCl, 20 mM Na_2_MoO_4_ and 2.5 mM desthiobiotin). RAF1-containing fractions were pooled, concentrated to <5 ml with a 30-kDa Vivaspin concentrator (Sartorius), and further purified by size exclusion chromatography (SEC) with a Superdex S200 16/600 column (GE Healthcare) in buffer A1, using an ÄKTA FPLC system (GE Healthcare). Protein standards [(Gel Filtration Calibration Kit (GE Healthcare)] were loaded onto the column for molecular weight calibration using the same method. Finally, the eluted peak corresponding to RAF1-CDC37-HSP90 complexes was collected and concentrated with a 30-kDa Vivaspin concentrator to final concentration of about 2 mg/ml and stored at −80°C until use.

### Cryo-EM sample preparation and data collection

For Cryo-EM, fresh RHC fractions from the Superdex S200, at a concentration of 0.6 mg/ml, were flash frozen in liquid nitrogen and stored at −80°C. Initially, the RHC samples were diluted with buffer A1 to a final concentration of 0.3 mg/ml for cryo-EM grid preparation, data collection and processing. Analysis of these initial results showed a strong preferential orientation of the RHC complexes in the grids. To alleviate the associated anisotropy in the obtained maps, additional tilted images (30°) were also collected as default protocol. Furthermore, we observed that the large heterogeneity of the RHC complexes, with a relatively small fraction of the particles yielding medium to high resolution maps, could be reduced when adding the 14-3-3 protein to the sample, improving the 3D classifications and reconstructions of the complex. Concentrated samples of 14-3-3ζ protein (5.3 mg/ml) were purified as previously reported (Lu et al., 2017). Final grids were prepared incubating the RHC complex (0.3 mg/ml final concentration in buffer A1) with the 14-3-3 protein (1:2.5 mole ratio, RHC:14-3-3ζ minutes in ice. A volume of 3 µl was applied to glow-discharged grids (Quantifoil R2/2 200 mesh Au grids; Leica EM ACE200, 50 s at 5 mA) and frozen using a Vitrobot Mark IV (FEI, Thermo Fisher Scientific; settings: 100% humidity, 277K and blot time 3 s). Grids were screened on a Glacios TEM (Thermo Fisher Scientific) and the best grid was transferred to a Titan Krios microscope (Thermo Fisher Scientific) operating at 300 kV at liquid nitrogen temperature. Dose-fractioned Movies (8138 in total: 4430 non-tilted and 3708 30° tilted; Figure S3a) were recorded using a Falcon 3EC Direct Electron Detector (Thermo Fisher Scientific) in electron counting mode (30 frames/Movie, with a dose rate of ∼1 e-/Å2 per frame, a defocus range of −0.5 to −2.5 µm and a pixel size of 0.832 Å/pixel).

### Cryo-EM data processing

The data from initial experiments were processed using cisTEM (Grant et al., 2018) and Relion (Zivanov et al., 2020), using in the preprocessing steeps MotionCor2 (Zheng et al., 2017) and CTFFIND4 (Rohou and Grigorieff, 2015). For the final data, obtained from the sample including 14-3-3ζ, all the preprocessing was performed in cryoSPARC (Punjani et al., 2017) because of the convenience of its patch motion correction and patch CTF estimation protocols for the analysis of medium-small particles and tilted images. A summary of the steps in the image processing procedures is shown in Figure S3c. Coordinates of the particles were determined with the blob picker protocol (cryoSPARC) and a wide range of sizes were selected to cover from the large RHC complexes to the smaller 14-3-3 dimers (Figure S3b). A total of 3.7 million particles were extracted (Figure S3c) and subjected to successive 2D and 3D classifications (cryoSPARC), using as initial volumes the 3D reconstructions obtained with the *Ab-initio* protocol (cryoSPARC and later also within Relion) applied to different subsets of 2D classes. Although the initial idea was to add 14-3-3 with the aim of isolate RHC:14-3-3 intermediate complexes, instead we discovered that only RAF1:14-3-3 complexes were obtained. Probably the binding of 14-3-3 to the CTD of RAF1 is too flexible and heterogeneous to be seen in the averaged 3D reconstructions, and maybe when the CTD peptides of RAF1 are fully accessible within the RHC complex the binding of 14-3-3 to them promotes the release of RAF1 from the HSP90 dimer. Accordingly, we classified the particles as RHC complexes (1.3 million particles), RAF1:14-3-3 complexes (1.2 million particles) and isolated 14-3-3 dimers (1.1 million particles). Although the processing was successfully continued inside cryoSPARC, extracted particles were imported, using pyEM (https://zenodo.org/record/3576630), into Relion for the final refinements and reconstructions. After 2D/3D classification and refinements, only medium-resolution maps, around 6 Å, were obtained from the isolated 14-3-3 particles possibly because of their flexibility, relatively low density, and small size (Figure S3c). Similarly, just medium to low-resolution maps were produced after classification and auto/manual refinements of the particles categorized as [RAF1 dimer + 14-3-3 dimer] (Figure S3c and S6). In those particles, the 14-3-3 part seems to be located at different distances to the RAF1 dimer and with different relative orientations because of the flexibility of the RAF1 interacting CT peptides. Even with additional classifications, signal subtraction (14-3-3 region), masking and local refinement of exclusively the RAF1 dimer, high-resolution maps cannot be obtained apparently because of the high heterogeneity of the C-lobes (that connect directly to the 14-3-3 dimer through their bound CT peptides). In the case of the RHC complex, a 33 % of the initial particles can be classified into a 3D class that, although still heterogeneous, can produce a high-resolution map after 3D auto-refinement (Relion; Figure S3c, 442k particles, 3.59 Å). The rest of the classes reflect a high degree of heterogeneity in all the components of the RHC complex, that prevents a proper refinement of the poses of the particles. All these classes could represent intermediate states of the RHC complex where the HSP90 dimer is transitioning to more open conformations or different rearrangements of their different domains. As we are obtaining RAF1:14-3-3 complexes, probably empty HSP90 particles could be attracted to these inconclusive 3D classes. Analysis of the variability of this 33% class using cryoSPARC(Punjani et al., 2017) and cryoDRGN (Zhong et al., 2021) showed that a large degree of flexibility, which is clearly localized at the RAF1 C-lobe and the visible regions of CDC37 (Figure 3b) and that those two lateral blobs of the RHC complex behave independently, not in a concerted way. Three different approaches were followed to classify and characterize the RHC particles. In the first one, an additional 3D classification of the 33% class was performed trying to achieve exclusively higher resolution. Hence, a map at 3.16 Å was obtained from 266k particles (RHC-I; Figure 1e, Figures S3c and S4a-e). Local high resolution was achieved at this map, but clearly limited to the HSP90 dimer and the RAF1extended segment that goes through it (Figure S4a-b); the corresponding regions to RAF1C-lobe and CDC37 MD showed lower resolution and visibly displayed the inherent not resolved heterogeneity, being not well defined. The use of tilted images helped in achieving a better distribution of orientations (Figures S3b and S4c-d) and reducing the anisotropy of the map (Figure S4e). In a second approach, a different set of classifications of the initial particles (Relion), now more focused in the characterization of the heterogeneity of RHC complex than trying to reach high-resolution, yielded eight different maps (Figure 3a and S3c). One of them, RHC-II shows the best defined RAF1 C-lobe and CDC37 MD, the latter containing an interacting portion of the RAF1 N-lobe (Figure 1f-g, Figure 3a and Figure S3c). This map presents similar characteristics than RHC-I, although the lower number of particles affect marginally its quality (local resolution, orientation distribution and anisotropy; Figure S4f-j). Other maps show different orientations of those two domains or worse defined densities, specially at the CDC37 MD region. Particularly interesting are the maps RHC-VI and RHC-VII (Figure 3a). The former looks like an empty closed HSP90 dimer but distinctively it shows densities corresponding with the RAF1 unfolded and extended segment (Figure 3a inset), therefore this complex should contain at least RAF1, although it seems to be freely moving and not attached to CDC37 ND (which is not visible). The peculiarity of map RHC-VII is that it apparently shows the CDC37 MD without its HSP90-anchored ND, so it should be interacting with the unfolded RAF1 N-lobe, although the RAF1 C-lobe is not visible due to its flexible attachment to HSP90 (free without CDC37 ND). These maps could represent transient conformations waiting for the release and subsequent proper folding of the RAF1 N-lobe. In the third approach, both lateral densities of the RHC complex were independently analyzed through masking and local classification and refinement (Relion; Figure 3c and Figure S3c). Ten different classes were obtained when focusing the classification on the RAF1 C-lobe, corresponding those clearly to slightly different rotations of that domain around a fulcrum, constituted by two close elements: the entry point of the extended segment into the HSP90 main body and the interaction of that C-lobe with the CDC37 ND (Figure 3c upper panel and 3d). Another ten classes were obtained when the focus was put on the other lateral blob, but in this case the maps were not as clear as in the case of RHC-II, so not clear details could be extracted about the state of the CDC37 MD and/or RAF1 N-lobe (Figure 3c lower panel). Final maps were refined, sharpened and their local resolution estimated in Relion; global resolution was estimated based on the gold-standard Fourier shell correlation 0.143 criterion.

### Model building and refinement

Previous models of the HSP90-CDC37-Cdk4 complex (PDB ID: 5FWK), B-Raf:14-3-3 complex (6UAN), CDC37 protein (1US7.) and RAF1 protein (3OMV) were used for the initial interpretation of the maps and preliminary model building. Also, we used Alphafold v2.0 (Tunyasuvunakool et al., 2021) predictions of RAF1 and CDC37. Both the CDC37 CTD and the N-terminal part of RAF1(∼349 aa) were not visible in the cryo-EM maps and probably they are moving freely without additional contacts with the rest of the RHC complex. Initial fitting of the models into the maps was performed with ChimeraX(Goddard et al., 2018). Coot(Emsley et al., 2010) was used for model building, modification, manual local refinement and visualization. Coordinate refinements in real space were performed with Phenix(Adams et al., 2010). Isolde (Croll, 2018)and Namdinator(Kidmose et al., 2019) were used for additional refinements in regions of low-resolution. Figures and videos were generated in ChimeraX and PyMOL https://pymol.org/2/.

### Western blot analysis and antibodies

For Western blotting, 30 μg of input and 20 μl of Strep pull-down eluate coming from Expi293F cells were separated by SDS-PAGE and transferred to nitrocellulose membranes (GE Healthcare). Membranes were probed with antibodies for HSP90 (Santa Cruz sc-13119, 1:1000), RAF1 (BD 610151, 1:1000), CDC37 (Santa Cruz sc-13129, 1:1000), 14-3-3 ε (Santa cruz sc-393177, 1:1000), 14-3-3 ζ (Santa cruz, sc-293415, 1:1000), A-RAF (Cell Signaling 4432, 1:500), B-RAF (Santa Cruz, sc-5284 1:1000), pan-Ras (Calbiochem OP40, 1:250), and Vinculin (Sigma V9131, 1:5000). Primary antibodies were detected with appropriate secondary antibodies conjugated to HRP and visualized by ECL-Plus (GE Healthcare).

### Protein Kinase assay

*In vitro* protein kinase assay was performed after Strep purification of RHC complex expressed in Expi293F cells. 2μg of the eluate was used for the kinase reaction, using a commercial RAF1 kinase assay kit (Millipore), following the protocol instructions of the provider. This assay kit is designed to measure RAF1 dependent phosphotransferase activity in a kinase reaction using recombinant inactive MEK1 as a RAF1 substrate. Samples were analysed by SDS-PAGE and the phosphorylation of MEK1 at residues Ser217/221 was detected by Western blotting with a commercial antibody (Cell Signalling #9121, 1:500).

### Sample preparation for Mass Spectrometry and Size-exclusion chromatography with multiangle light scattering (SEC-MALS) analysis

For MS-based analysis of RHC complex phosphorylation status, the proteins were resolved by SDS-PAGE, using 4–12% bis-tris gradient gels (Invitrogen), and stained with the Colloidal Blue Kit (Invitrogen) according to manufacturer instructions. For each sample, the RAF1, HSP90 and CDC37 specific bands were excised from the gel in 1 × 2-mm cubes. Excised SDS-PAGE bands were washed, and proteins digested using the standard in-gel digestion procedure. Briefly, proteins were reduced (25 mM TCEP, 30 min. 45 °C) and alkylated (40 mM CAA, 1 h, RT in the dark) and digested with trypsin for 16 hours at 37 °C in 50 mM NaHCO_3_ (Promega) (estimated protein:enzyme ratio 1:100). Digestion was quenched with 0.1% TFA and resulting peptides were desalted using C_18_ stage-tips. In the case of proteins in solution, samples were digested by means of standard FASP protocol. Briefly, proteins were alkylated (50 mM CAA, 20 min in the dark, RT) and sequentially digested with Lys-C (Wako) (estimated protein:enzyme ratio 1:100, o/n at RT) and trypsin (Promega) (estimated protein:enzyme ratio 1:100, 6 hours at 37°C). Resulting peptides were desalted using C_18_ stage-tips.

LC-MS/MS was carried out by coupling an UltiMate 3000 RSLCnano LC system to either a Q Exactive Plus or Q Exactive HF mass spectrometer (Thermo Fisher Scientific). In both cases, peptides were loaded into a trap column (Acclaim™ PepMap™ 100 C18 LC Columns 5 µm, 20 mm length) for 3 min at a flow rate of 10 µl/min in 0.1% FA. Then, peptides were transferred to an EASY-Spray PepMap RSLC C18 column (Thermo) (2 µm, 75 µm x 50 cm) operated at 45 °C and separated using a 60 min effective gradient (buffer A: 0.1% FA; buffer B: 100% ACN, 0.1% FA) at a flow rate of 250 nL/min. The gradient used was, from 2% to 6% of buffer B in 2 min, from 6% to 33% B in 58 minutes, from 33% to 45% in 2 minutes, plus 10 additional minutes at 98% B. Peptides were sprayed at 1.5 kV into the mass spectrometer via the EASY-Spray source and the capillary temperature was set to 300 °C.

The Q Exactive Plus mass spectrometer was operated in a data-dependent mode, with an automatic switch between MS and MS/MS scans using a top 15 method. (Intensity threshold ≥ 4.5e4, dynamic exclusion of 10 sec and excluding charges unassigned, +1 and > +6). MS spectra were acquired from 350 to 1500 m/z with a resolution of 70,000 FWHM (200 m/z). Ion peptides were isolated using a 2.0 Th window and fragmented using higher-energy collisional dissociation (HCD) with a normalized collision energy of 27. MS/MS spectra resolution was set to 35,000 (200 m/z). The ion target values were 3e6 for MS (maximum IT of 25 ms) and 1e5 for MS/MS (maximum IT of 110 msec).

The Q Exactive HF mass spectrometer was operated as described above for the Q Exactive Plus mass spectrometer except using an Intensity threshold ≥ 2.2.e4, dynamic exclusion of 10 sec and excluding charges +1 and > +6. MS spectra were acquired from 350 to 1400 m/z with a resolution of 60,000 FWHM (200 m/z). Ion peptides were isolated using a 2.0 Th window and fragmented using higher-energy collisional dissociation (HCD) with a normalized collision energy of 27. MS/MS spectra resolution was set to 30,000 (200 m/z). The ion target values were 3e6 for MS (maximum IT of 25 ms) and 1e5 for MS/MS (maximum IT of 45 msec).

### Size-exclusion chromatography with multiangle light scattering (SEC-MALS) analysis

The oligomeric state and stoichiometry of the RHC complex were analysed by SEC-MALS analysis. Proteins samples were prepared as described previously. 105 µg of sample (at 0.7 mg/mL) were injected in a Superdex 200 10/300 size exclusion column (GE Healthcare) equilibrated in buffer A1. Data were collected on Dawn Heleos 8+ and Optilab TRex detectors and analyzed using ASTRA6 software (Wyatt Technology). The monomeric BSA (Sigma-Aldrich) was used as standard to verify instrument performance.

### CDC37 purification and binding assays on cellulose-bound peptides containing RAF family sequences

The human full-length CDC37 cDNA was amplified by PCR with restriction site-tailed primers incorporating a C-terminal StrepTagII, separated from protein by a Gly-Ser-Ala residue linker from the Origene RC201002 plasmid. The resulting DNA was cloned into the HindIII/XhoI sites of the transfer vector pcDNA3 (Invitrogen), using the In-Fusion® HD EcoDry Cloning Kit (Takara Bio USA). The derivative plasmid, pcDNA3-CDC37-Strep, was sequence verified. Four 120 ml cultures of Expi293F cells were transfected with 120 µg of pcDNA3-CDC37-Strep plasmid using the ExpiFectamine 293 Transfection Kit as described for the RHC complex. After the first step of protein purification by affinity chromatography, CDC37-containing fractions were pooled, concentrated to <5 ml with a 10-kDa Vivaspin concentrator (Sartorius), and further purified by size exclusion chromatography (SEC) with a Superdex S75 16/600 column (GE Healthcare) in buffer A1, using an ÄKTA FPLC system (GE Healthcare). The eluted peak corresponding to CDC37 protein was collected and concentrated with a 10-kDa Vivaspin concentrator to final concentration of about 2 mg/ml and stored at −80 °C until use.

Overlapping dodecapeptides scanning the kinase domain sequence of human A-RAF, B-RAF and RAF1 were prepared by SPOT-Synthesis (JPT, Berlin, Germany) onto a cellulose-βalanine-membrane (Table S3). The membranes were rinsed with methanol, washed with TBS-T and blocked for 2 hours using 2.5% BSA. The saturated membranes were incubated with purified CDC37 protein (10μg/ml) for 3 hours and after several washing steps with TBS-T, incubated with anti-CDC37 antibody (Santa Cruz Biotechnology), followed by HRP-conjugated secondary antibody (Dako). Positive spots were visualized using the ECL system (Bio-Rad). Dots from the scanned films were quantified using the software ImageJ. To compare among the three RAF family members, a peptide with a common sequence was selected for normalization (peptide 18 from region #3). CDC37 binding affinity to each peptide was calculated as the mean pixel integrated density of each dot relative to a reference one.

### Site-directed mutagenesis and pull-downs from cell lysates

All RAF1 and CDC37 constructs expressed in mammalian cells for Figure 2 were generated by QuickChange Lightning Site-Directed Mutagenesis (Agilent Technologies), using the primer sequences shown in Table S4. RAF1 mutants including L459E, I464E, K483A, I484A, D486A, F487A, F487R and F487E were generated from pcDNA3-RAF1-Strep plasmid. Human RAF1-CAAX cDNAs carrying the D486A, F487A and D486A/F487A mutations used for retroviral expression in K-Raslox MEFs were generated from pBABE-puro-Raf1-WT-CAAX plasmid. The CDC37 I23E mutant was generated from pcDNA3-CDC37-Strep plasmid.

For the generation of RHC complexes carrying the above RAF1 mutants, 20 ml cultures of Expi293F cells were transiently transfected with 20 μ g pcDNA3-RAF1-Strep wt or mutant variant + 5 μ pcDNA3-HSP90-HA + 5 µg of pCMV6-CDC37-Myc-DDK) using polyethylenimine (PEI) (Polysciences Inc.). Cells were harvested 48 h after transfection. Pellets were lysed as described before. 20 mg of total protein extracts were incubated with 200 μl pre-washed Strep-Tactin resin (IBA GmbH) for 1 hour at 4 °C. Resin was spun down (1000×g for 30 s) and washed five times with 500 μl buffer A1. Proteins were eluted from the Strep-Tactin matrix with 2×100μl of buffer A1 supplemented with 5 mM desthiobiotin for 10 min on ice. For the CDC37 interaction mutant experiment, 20 ml cultures of Expi293F cells were transiently transfected with 20 μg plasmid DNA (10 μg pcDNA3-RAF1-His + 5 μg HSP90-HA + 5 µg of pcDNA3-CDC37-Strep wt or I23E) using PEI and processed as described for RAF1 mutants. In both cases, eluates were analysed on SDS-PAGE, followed by Western blotting with specific antibodies and staining with Coomassie blue. To get an accurate quantification of the proteins, part of the eluate was analysed by Mass Spectrometry.

### Processing and analysis of MS data

Raw files were processed with MaxQuant using the standard settings against a human protein database (Swiss-Prot, 20,373 sequences) containing the recombinant protein sequences used in this work and supplemented with contaminants. Carbamidomethylation of cysteines was set as a fixed modification whereas oxidation of methionines, protein N-term acetylation and phosphorylation of serines, threnonines and tyrosines were set as variable modifications. Minimal peptide length was set to 7 amino acids and a maximum of two tryptic missed cleavages were allowed. Results were filtered at 1% FDR (peptide and protein level).

For RAF1 pull down experiments, the normalization factor was calculated by adding the intensities of all the RAF1 peptides whose sequences are shared by all RAF1 mutants, excluding those sequences in which a phosphorylation site was identified. For the CDC37 pull down, the normalization factor was calculated by summing the intensities of all the CDC37 peptides whose sequences are shared by all CDC37 mutants.

### Colony assay in *Rasless* and *Rafless* MEFs

Low passage (p12-15), immortal *H-Ras^-/-^;N-Ras^-/-^;K-Ras^lox/lox^* (K-Raslox) and *A-Raf^lox/lox^*;*B-Raf^lox/lox^*;*Raf1^lox/lox^* (Raflox) MEFs were infected with retroviral and lentiviral supernatants, respectively, and selected with the appropriate antibiotic as described earlier (Martín et al., 2005). Resistant cells were seeded in equal cell number (3,000-6,000) in the absence or presence of 4-hydroxytamoxifen (4OHT, Sigma, 600 nM). Cells were allowed to form colonies for 14 days. Plates were fixed with 1% glutaraldehyde (Sigma), stained with crystal violet (Merk), and colonies > 2 mm in diameter scored. When needed, representative colonies were picked and expanded for further analysis. In addition to the indicated cDNAs, all K-Raslox cells were infected with a retrovirus encoding a shRNA directed against p16INK4a-specific sequences, to prevent cellular senescence (Drosten et al., 2010). This step was skipped in Raflox cells.

### Peptides for exogenous delivery and cell proliferation assays

Peptides corresponding to sequences M456 to D468 and I484 to W496 of RAF1 and an HA control peptide were fused to an N-terminal cell penetrating motif (TAT motif) via a (GlySer)_3_ linker sequence and chemically synthesized by GenScript’s Custom Peptide Synthesis service at crude purity. For dose-response experiments, the human lung tumor cell line A549 and the human patient derived xenograft (PDX) Pdc1 were seeded in 96 well plates (n=4) at 50% confluency and peptides were added at the indicated concentrations. Cell viability was determined after 72 hours using a colorimetric assay based on 3-(4,5-dimethylthiazol-2-yl)-2,5 diphenyltetrazolium bromide (MTT) metabolization. Cell viability was normalized to that of an untreated control on the same plate. The experiment was repeated 3 times.

## SUPPLEMENTAL INFORMATION

**Figure S1. Related to Figure 1 and 4. Structural alignment of the kinase domains of the RAF family of proteins.** Amino acid sequences of human RAF1, A-RAF and B-RAF proteins were aligned by Clustal Omega (http://www.ebi.ac.uk/Tools/msa/clustalo). The Figure was prepared with ESPript (http://espript.ibcp.fr). Residue numbers are labelled according to the RAF1 sequence. Similar residues are shown in red and identical residues in white over red background. Secondary structures including α helix and β sheets for RAF1 and *B-*RAF are indicated. The violet rectangle includes the unfolded segment of RAF1 visualized in the RHC complex. The green rectangle shows the disordered region of RAF1 observed in the crystal structure of the kinase domain.

**Figure S2. Related to Figure 1. Purification and characterization of the RAF1-HSP90-CDC37 complex used for cryo-EM structure determination.** a) Elution profile from size exclusion chromatography (SEC) on Superdex 200 column. b) Coomassie-stained SDS-PAGE analysis of eluted fractions. c) SEC-MALS profile of RHC purification. The chromatogram shows a major broad peak whose central part is homogeneous and corresponds to a molecular weight of 268 kDa (265-272 kDa) indicating a stoichiometry of 1:2:1 RAF1:HSP90:CDC37 (^Theor.^M_w_ = 290 kDa, 74+2×84+48). This peak also shows a heterogeneous left shoulder compatible with the presence of larger species (270-325 kDa) that could correspond to other RHC complexes with different stoichiometry. The small peak on the right of the chromatogram (71-89 kDa) corresponds primarily to HSP70, as determined by mass spectrometry. d) Mass spectrometry analysis of the affinity-purified extract containing the RHC complex as well as co-purified 14-3-3 proteins. The plot shows the mean protein levels of three different experiments, normalized to RAF1 levels. e) Coomassie-stained SDS-PAGE analysis of purified 14-3-3ζ.

**Figure S3. Related to Figures 1, 2, 3 and S6. Cryo-EM image processing. a)** A representative motion-corrected electron micrograph of the (RHC + 14-3-3ζ mixture embedded in vitreous ice on a gold grid at −2 µm defocus. **b)** Representative reference-free 2D class averages of particle images showing secondary structure elements (box dimensions: 384×384 pixel^2^, 319×319 Å^2^; cryoSPARC). Projections of the RHC complex can be easily identified and its different subunits are indicated (H - HSP90, R - RAF1, C - CDC37). **c)** Summary of the cryo-EM data processing workflow showing the different procedures used to analyze the different particles present in the sample: RHC complexes, 14-3-3 dimers and RAF1:14-3-3 complexes. The number of particles and the global resolution of certain classes are indicated.

**Figure S4. Related to Figures 1, 2 and 3. Analysis and validation of RHC-I and RHC-II maps. a-e)** RHC-I complex**. f-j)** RHC-II complex**. a, f)** Local resolution filtered density maps coloured by local resolution (estimated with Relion). **b, g)** Histograms of the distribution of the local resolution estimation (Relion). **c, h)** Visual representations of the distribution of particles according to the angular sampling (Relion; the length and color of the cylinders correspond with the number of particles). **d, i)** Alternative depictions of the distribution of available orientations (Cryosparc): (**upper panel**) number of images per orientation and (**lower panel**) number of measurements per radial wedge. **e, j)** FSCs and analysis of the anisotropy of the two maps (calculated with 3DFSC).

**Figure S5. Related to Figures 1, 2 and 3. Detailed view of the interactions between RAF1 and the HSP90-CDC37 chaperone system. a)** Superposition of RAF1 kinase domain structure (white) and RAF1 in the RHC complex (green). The pink and magenta segments correspond to the section of RAF1 which folds into β4, β5, the connecting loops and the first turn of the αC helix in the kinase domain structure but is unfolded in the RHC complex. **b)** Section of RAF1 inside the lumen of HSP90. The chaperone structure is depicted with a surface representation colored according to the electrostatic potential while the cryo-EM map is shown on the RAF1 fragment. **c)** Region of RAF1 interacting with the MD/CTD region of CDC37. The surface representation of CDC37 is colored according to its electrostatic potential, while the RAF1 region is overlayed with the cryo-EM map. **d)** Scheme depicting the interaction sections between CDC37 and RAF1 in the RHC complex. Fitting of the model to the map in the interaction stabilizing the unfolded RAF1 including the region between the T19-F29 loop of CDC37 and the C-lobe of RAF1. **e)** Electrostatic potential map of the RAF1 C-lobe in the interacting region with the stabilizing loop of CDC37. **f)** Detailed view of the cryo-EM map fitting in the S13 phosphorylation showing the network of contacts facilitating the CDC37 conformation.

**Figure S6. Related to Figure 1 and Table S1. Conformational heterogeneity of the RAF1:14-3-3 complex. a)** Eight different 3D classes obtained from the initial pool of RAF1:14-3-3 particles. Additional refinement of one of the classes (blue) provided an improved map**. b)** Fitting of the models of the components of the RAF1:14-3-3 complex into the best map (RAF1, lime and green; 14-3-3 blue and purple).

**Figure S7. Related to Figure 4. Effect of a peptide mimicking the RAF1 region of interaction with CDC37 on lung tumor cell lines. a)** Diagram depicting the assay to validate the top binding peptide (RAF1 Pep484) identified in Figure 4b-c conjugated to the TAT protein transduction motif. **b)** Cell proliferation assay of A549 human lung cancer cell line and cells derived from the Pdc1 PDX lung tumor treated with increasing doses of a TAT-HA peptide used as negative control, the low interaction peptide TAT-RAF1 Pep456 and the TAT-RAF1 Pep484 high interaction peptide.

**Figure S8. Related to Figure 4 and 6. Structural comparison of the kinase domains of RAF1, B-RAF and mutant B-RAF V600E**. Superposition of the kinase domains of RAF1, B-RAF and B-RAF V600E mutant highlights the structural differences between these kinase domains. The dashed oval focus in the L480-P505 residue region where RAF1 interacts with CDC37 in the RHC complex. The differences in that area between B-RAF and the B-RAF V600E mutant together with the similarity of the B-RAF V600E kinase domain with RAF1, suggest why this oncogenic mutant interacts with the HSP90-CDC37 chaperone system.

**Table S1. Related to Figure 1. Phosphorylation sites (see Excel File).**

**Table S2.** Related to Figure 1, S3 and S4. Cryo-EM data collection, refinement and validation statistics.

| Model | RHC-I | RHC-II |
| --- | --- | --- |
| PDB code | 7Z38 | 7Z37 |
| EMDB code | EMD-14473 | EMD-14472 |
| Data collection |  |  |
| Electron Microscope | Titan Krios |  |
| Voltage (kV) | 300 |  |
| Detector | Falcon III (counting mode) |  |
| Movies | 8138 (4430 non-tilted + 3708 tilted) |  |
| Pixel size (Å/pixel) | 0.832 |  |
| Electron dose (e<sup>-</sup>/Å<sup>2</sup>) | 30 |  |
| Defocus range (mm) | 0.5-2.5 |  |
| 3D Reconstruction |  |  |
| Main software | Relion 3.13, Cryosparc 3.3.1 |  |
| Global resolution (Å) | 3.16 | 3.67 |
| Local resolution (Å) | 3.03-4.87 | 3.49-7.29 |
| number of particles | 266k | 46k |
| Map-sharpening b factor (Å<sup>2</sup>) | -98 | -78 |
| Model fitting |  |  |
| Software | ChimeraX, Phenix |  |
| Refinement |  |  |
| Software | Phenix, Coot, Isolde, Namdinator, Alphafold |  |
| Correlation coefficient CC (mask)* | 0.82 | 0.82 |
| Model composition |  |  |
| Chains | 4 | 4 |
| Atoms | 14264 | 14262 |
| Residues | Protein: 1740 | Protein: 1740 |
| Ligands | Mg: 2 | Mg: 0 |
|  | ATP: 2 | ATP: 2 |
| Validation |  |  |
| MolProbity score † | 1.94 | 2.00 |
| Clash score † | 10.61 | 14.33 |
| Ramachandran plot (%) * |  |  |
| Outliers | 0.06 | 0.17 |
| Allowed | 5.80 | 4.64 |
| Favored | 94.14 | 95.18 |
| Rotamer outliers (%) * | 0.25 | 0.25 |
| Bonds (RMSD) * |  |  |
| Length (Å) | 0.005 | 0.008 |
| Angles (°) | 0.605 | 0.977 |
| * as reported by Phenix <sup>(Adams et al., 2011)</sup> |  |  |
| † as reported by Molprobit (Chen et al., 2010) |  |  |

**Table S3.**
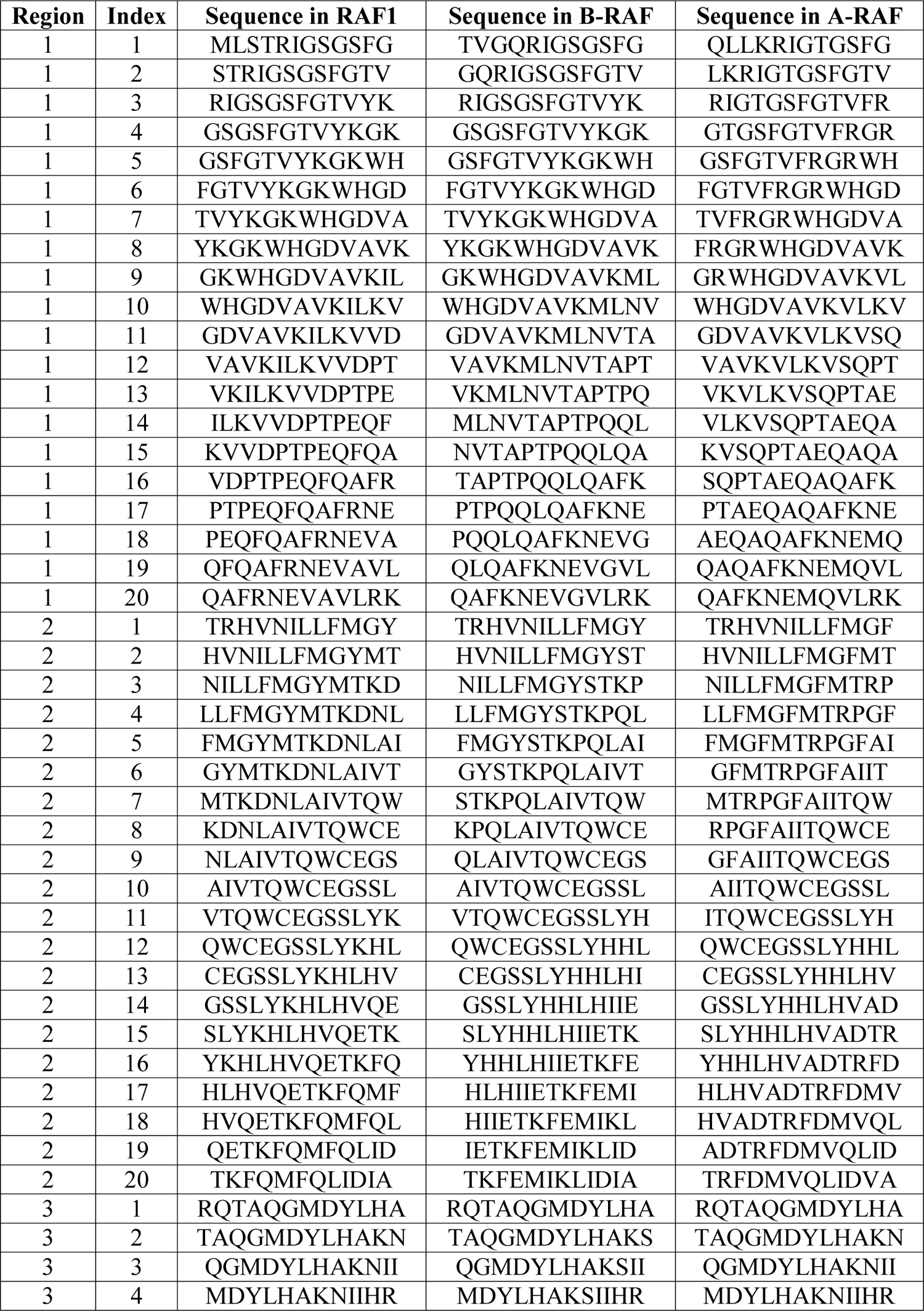
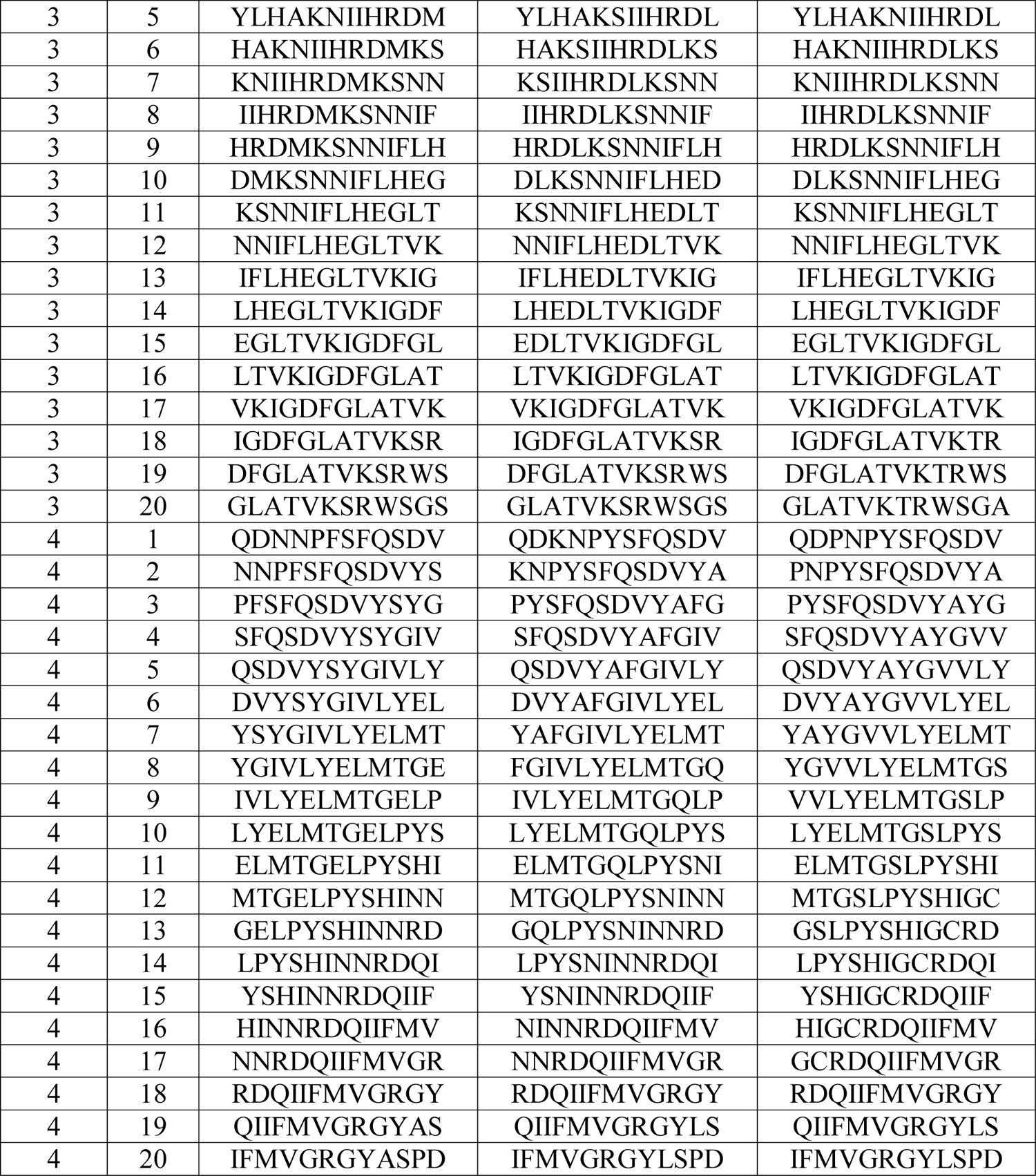
Related to Figures 4 and S7. Peptides used in the PepScan analysis.

**Table S4.**
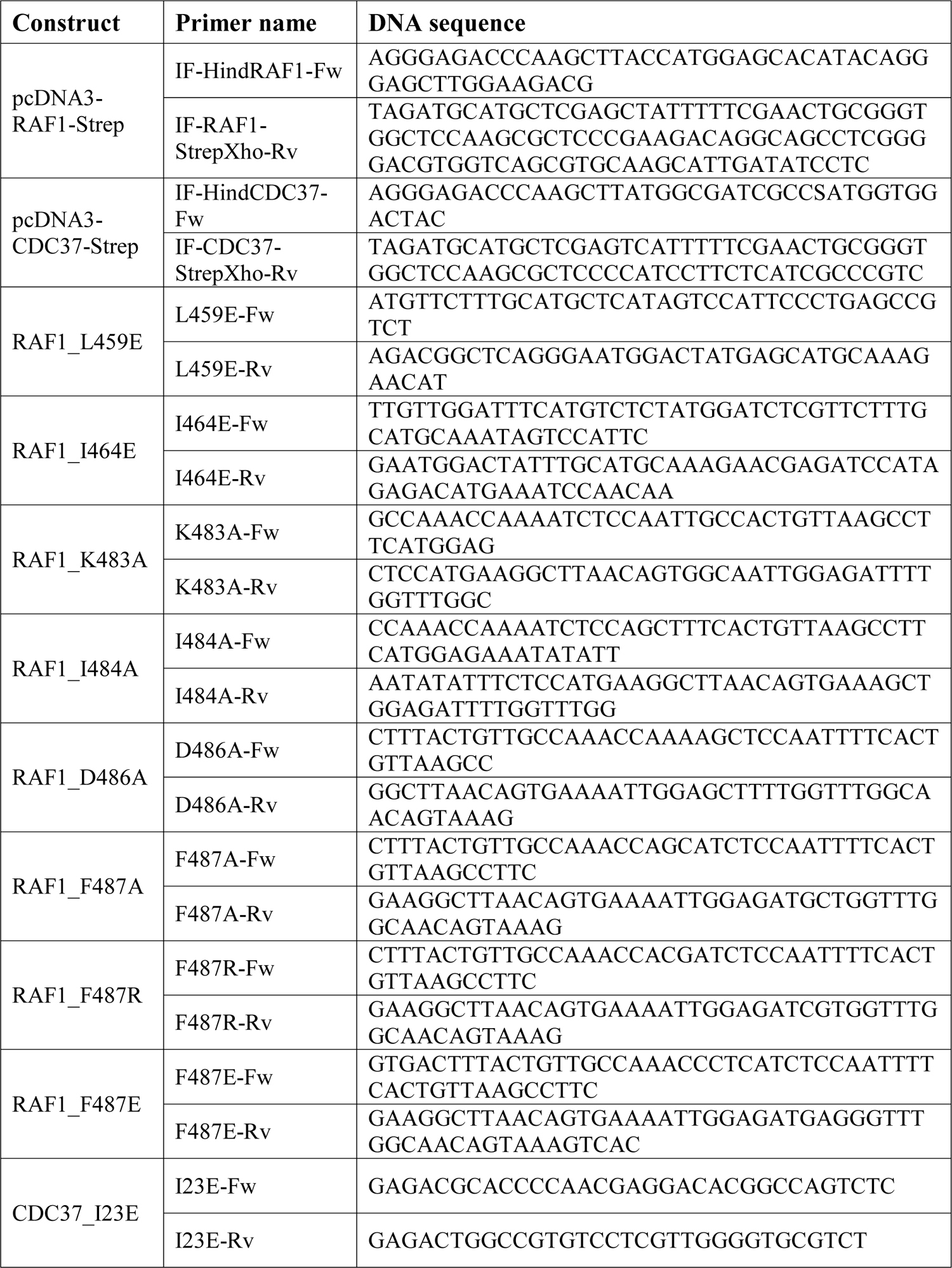
Related to STAR Methods. Primer sequences.

**Movie S1. Related to Figures 1 and 3.** 3D variability analysis showing the conformational heterogeneity of the RHC complex. The video displays the 3D variability analysis performed with cryoSPARC. Large variability is observed in the RAF1 C-lobe and CDC37 NTD and MD domains.

**Movie S2. Related to Figure 3.** Front view of the detailed view of the movements of the RAF1 C-lobe in the RHC complex in the region of interaction with CDC37. The loop comprising residues T19 to R32 in CDC37 engage in interactions with the activation loop of RAF1 mimicking the interactions with a folded RAF1 N-lobe to stabilize the C-lobe moiety.

**Movie S3. Related to Figure 3.** Back view of the movements of the RAF1 C-lobe in the RHC complex in the region of interaction with CDC37. The loop comprising residues T19 to R32 in CDC37 engage in interactions with the activation loop of RAF1 mimicking the interactions with a folded RAF1 N-lobe to stabilize the C-lobe moiety.

