## Supplementary material for "Structure of the RAF1-HSP90-CDC37 complex reveals the basis of RAF1 regulation": supp fig 1-8

### Slide 1
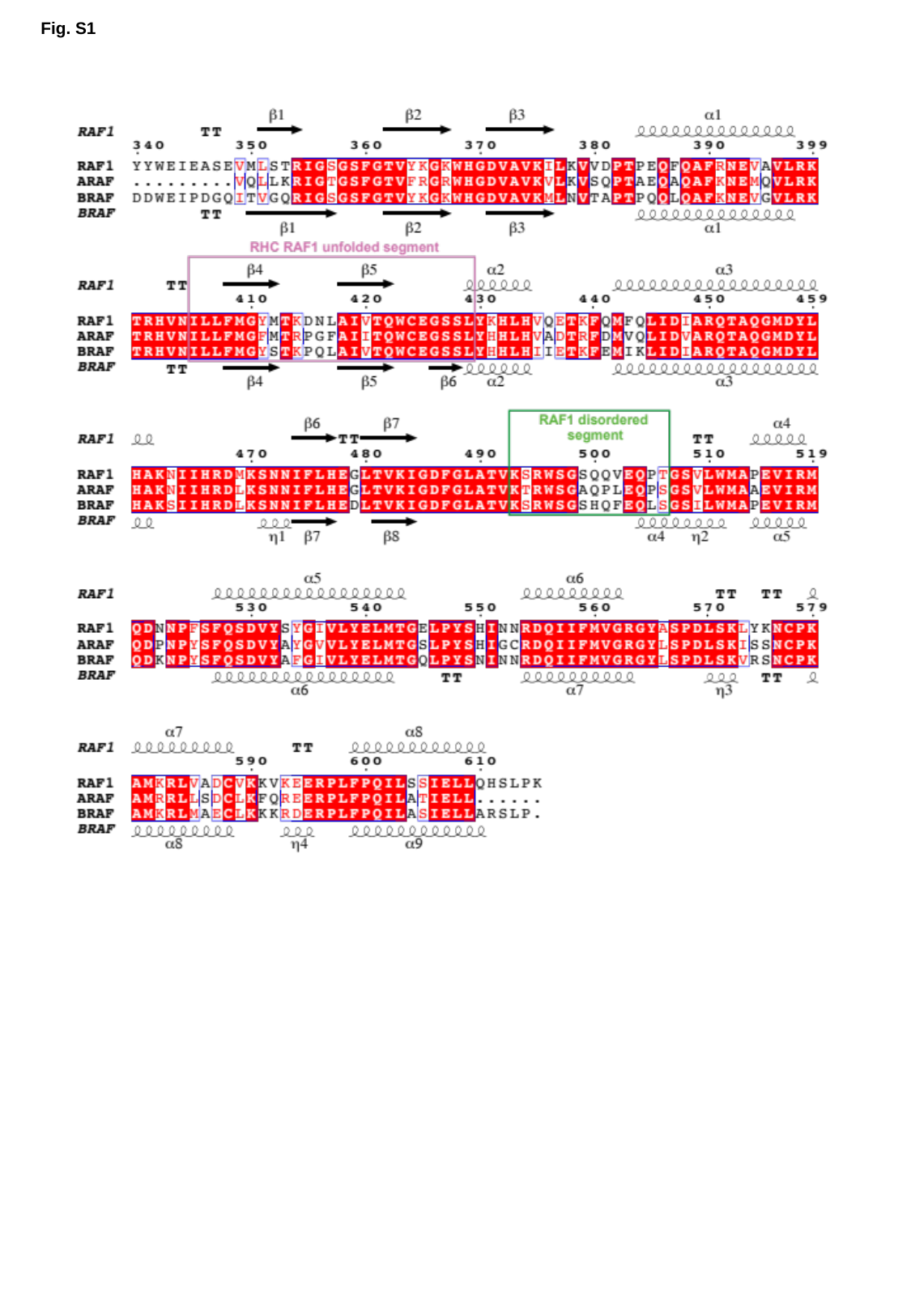

Fig. S1

### Slide 2
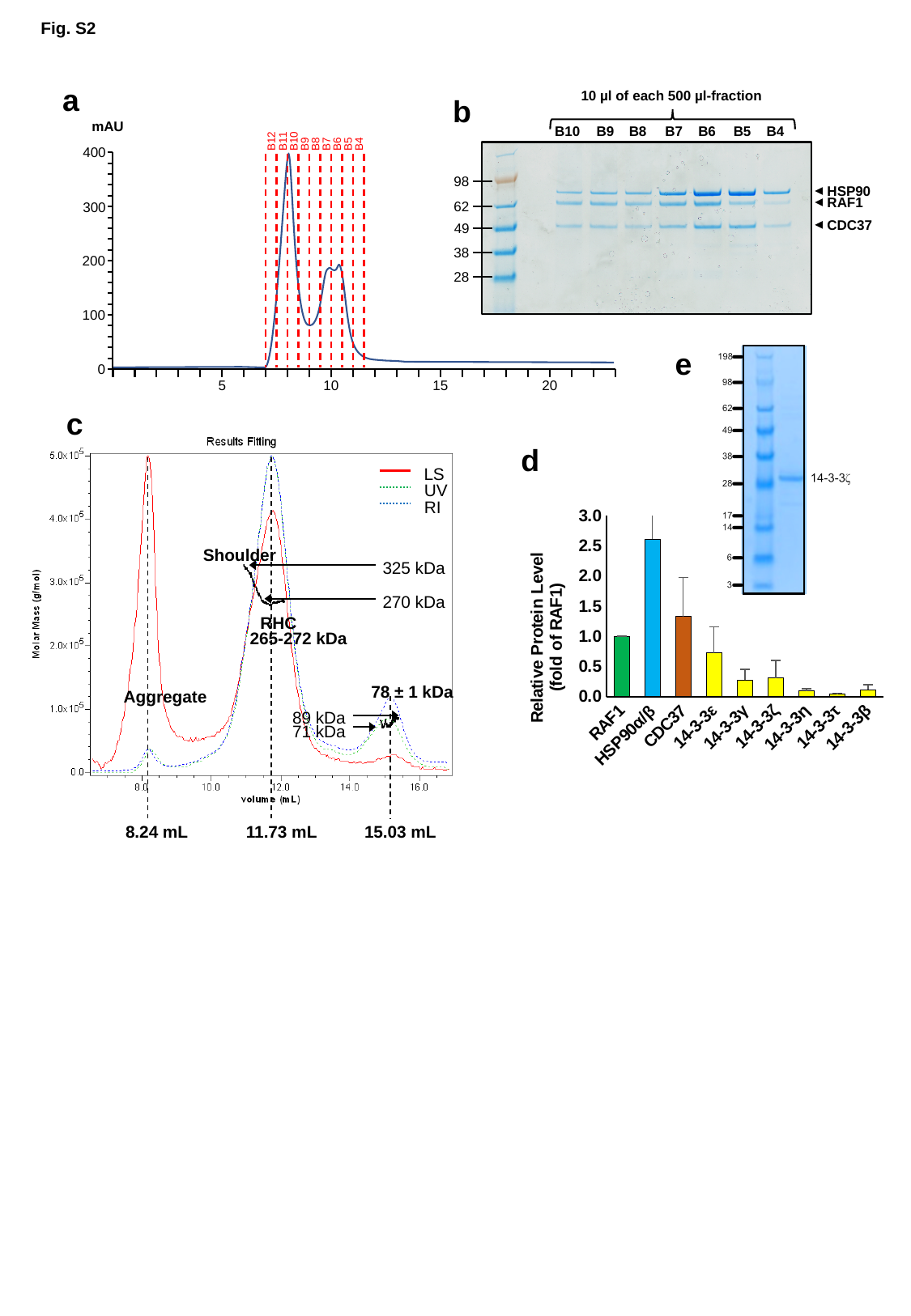

Fig. S2
a
b
10 µl of each 500 µl-fraction
B10
B9
B8
B7
B6
B5
B4
98
HSP90
RAF1
62
CDC37
49
38
28
mAU
B11
B10
B12
B9
B8
B7
B6
B5
B4
400
300
200
100
0
20
5
10
15
e
c
Shoulder
325 kDa
270 kDa
RHC
265-272 kDa
78 ± 1 kDa
Aggregate
89 kDa
71 kDa
8.24 mL
11.73 mL
15.03 mL
LS
UV
RI
d
#### Chart
| Category | |
|---|---|
| RAF1 | 1.0 |
| HSP90α/β | 2.606018618285835 |
| CDC37 | 1.3297503597291291 |
| 14-3-3ε | 0.7263932660680373 |
| 14-3-3γ | 0.27796611075487254 |
| 14-3-3ζ | 0.3098658989242443 |
| 14-3-3η | 0.09560965747927169 |
| 14-3-3τ | 0.04170311533278225 |
| 14-3-3β | 0.11407120422842654 |

### Slide 3
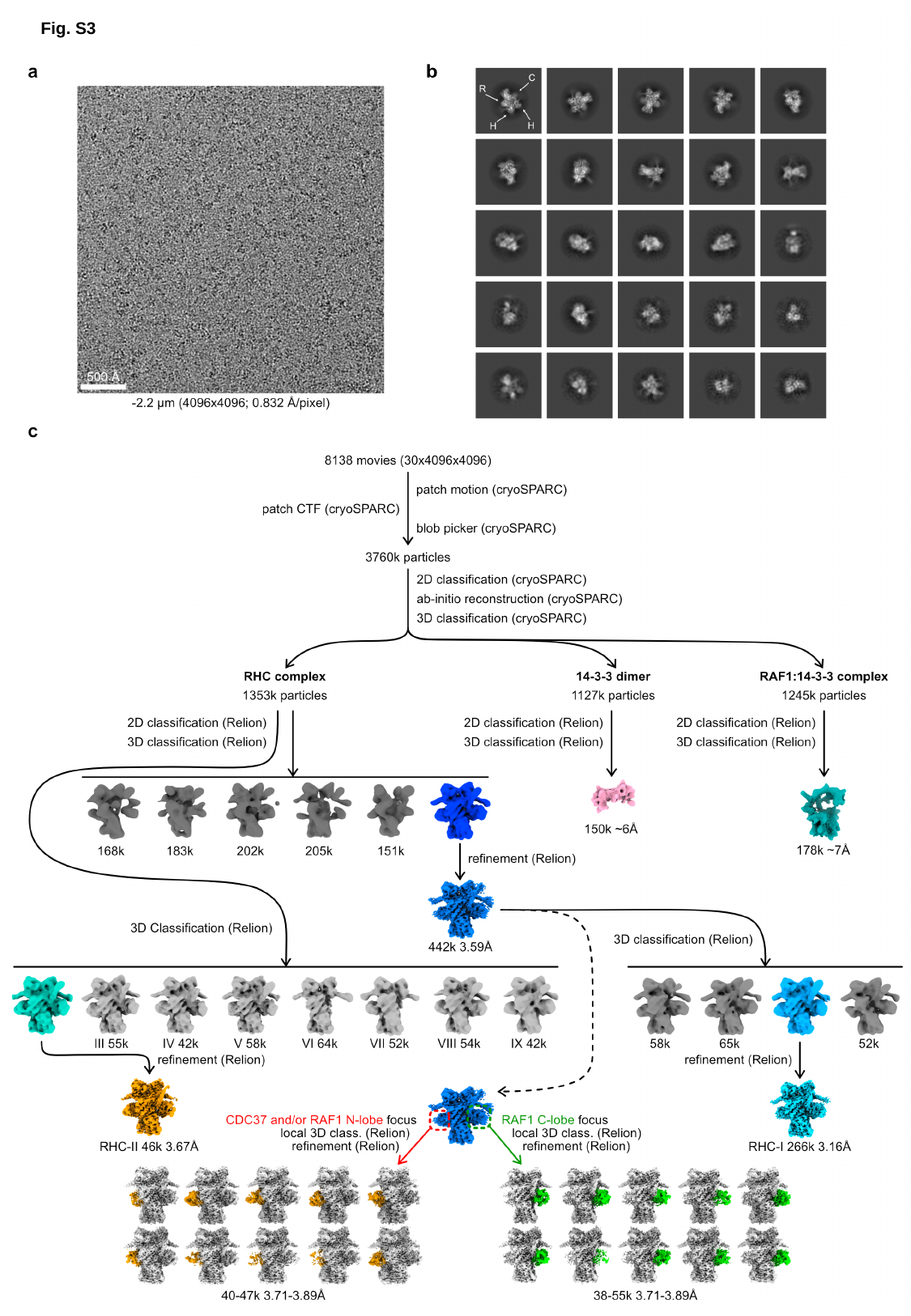

Fig. S3

### Slide 4
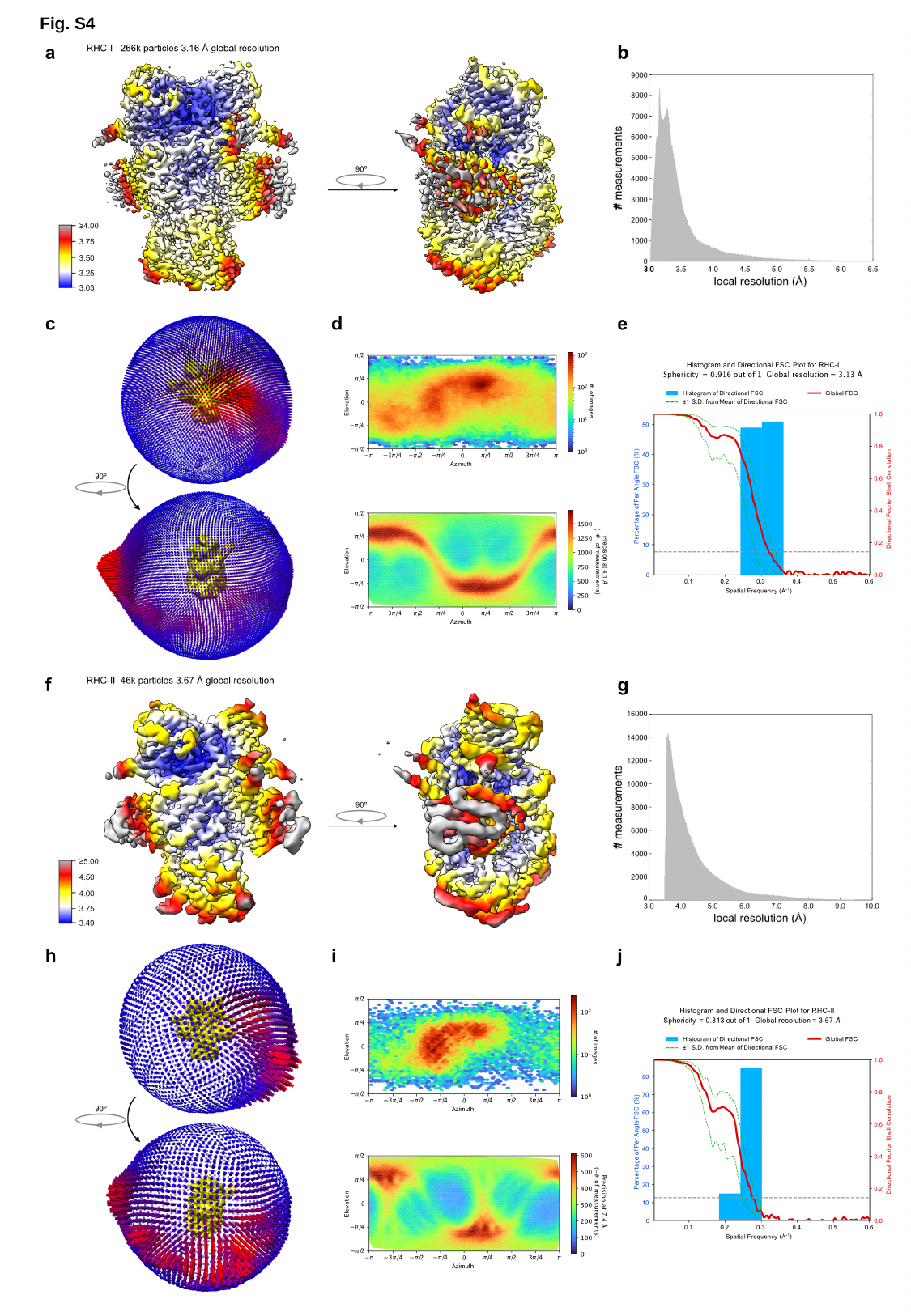

Fig. S4
Extended Data Fig. 4
Extended Data Fig. 4

### Slide 5
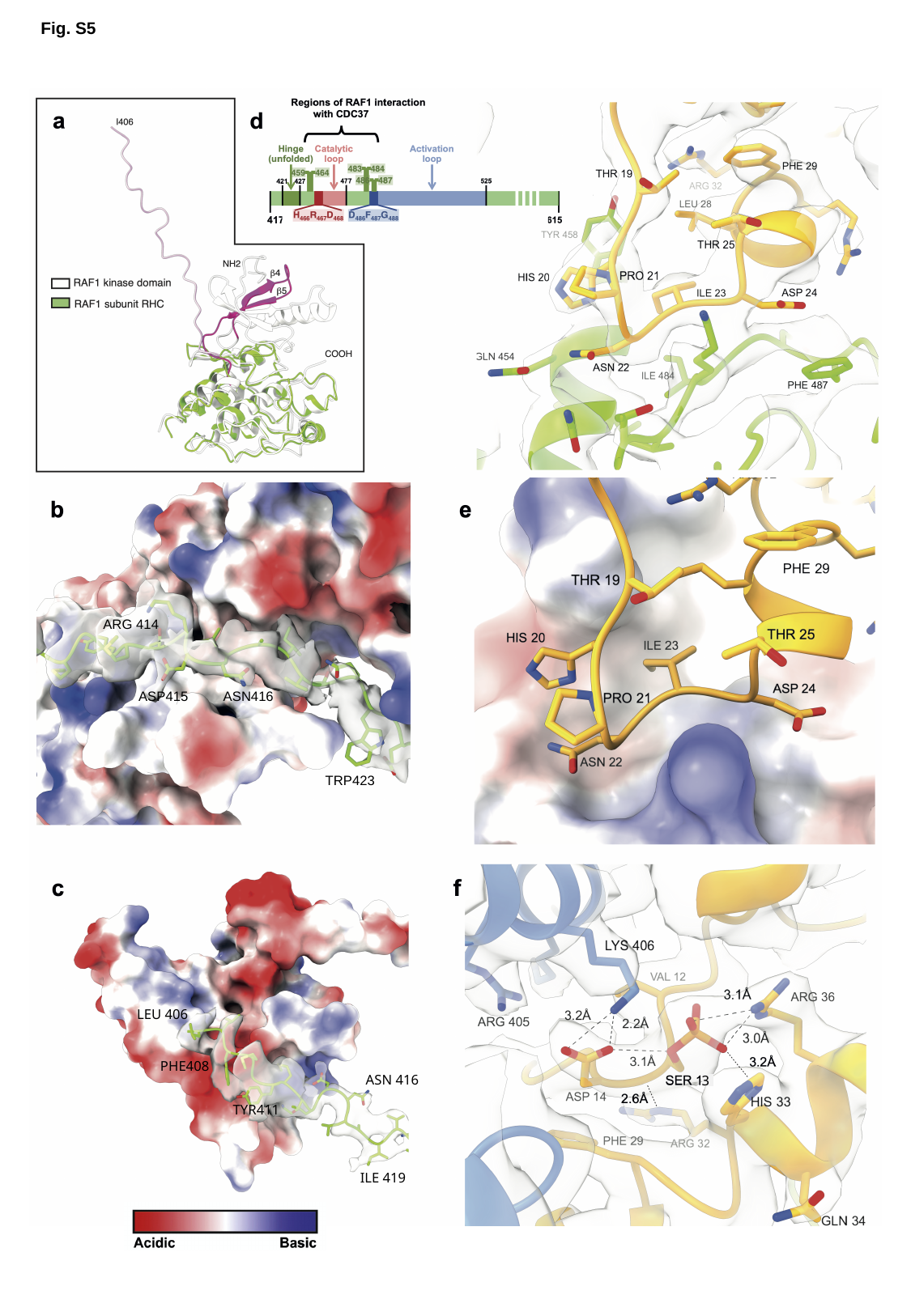

Fig. S5
ARG 414
ASP415
ASN416
TRP423
LEU 406
PHE408
ASN 416
3
TYR411
ILE 419

### Slide 6
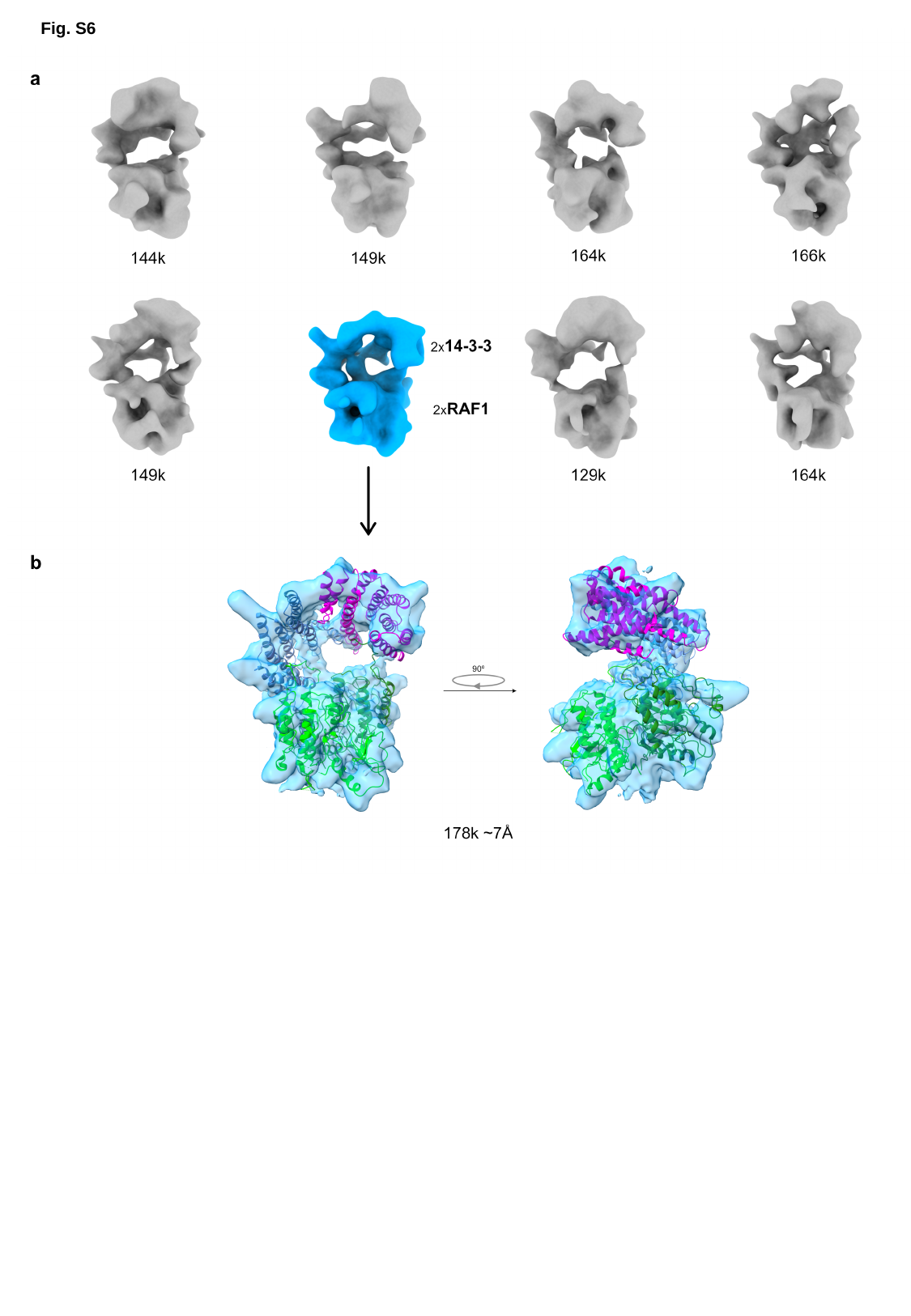

Fig. S6

### Slide 7
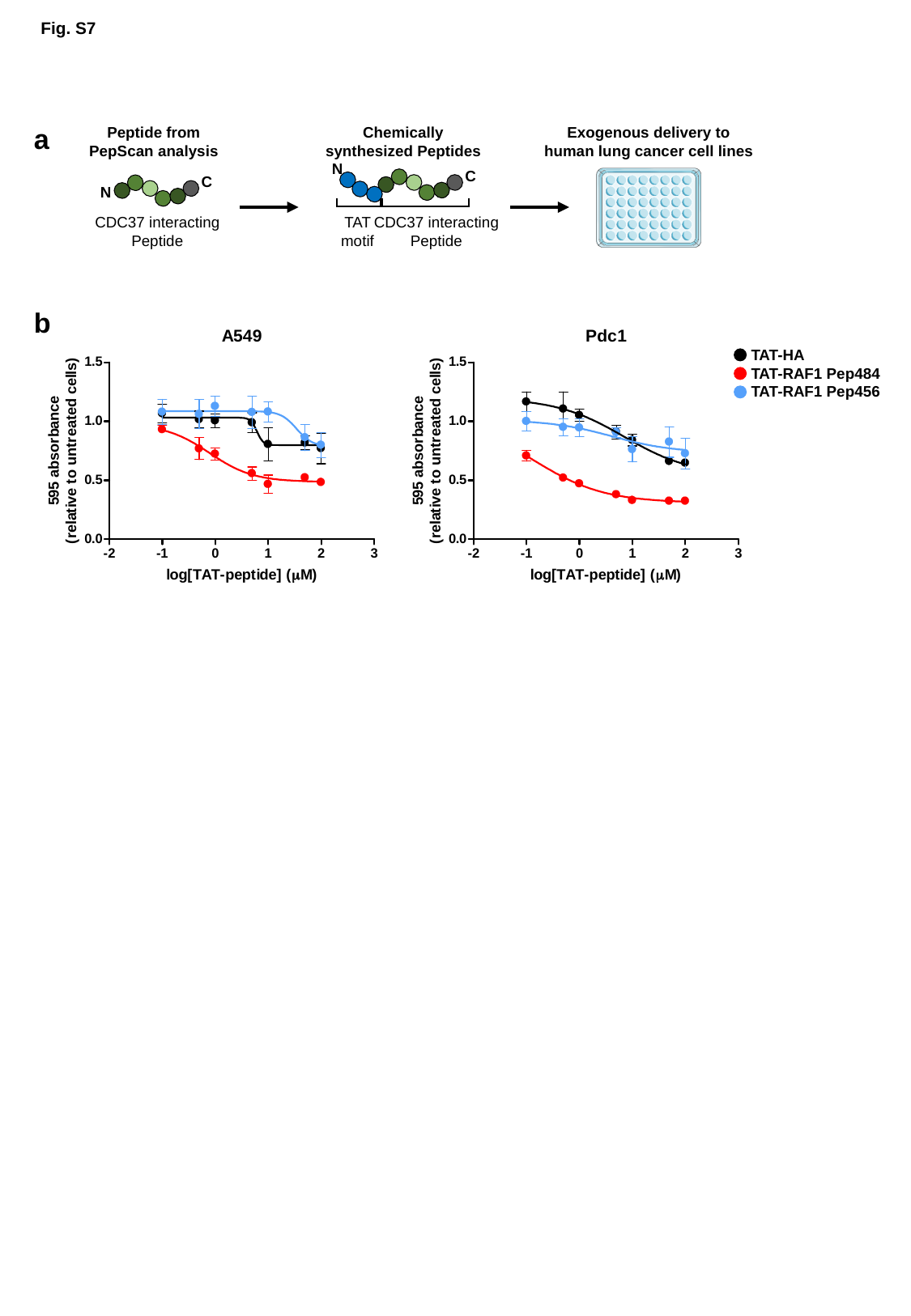

Fig. S7
a
Peptide from
PepScan analysis
Chemically synthesized Peptides
Exogenous delivery to
human lung cancer cell lines
N
C
CDC37 interacting Peptide
TAT
motif
C
N
CDC37 interacting Peptide
b
TAT-HA
TAT-RAF1 Pep484
TAT-RAF1 Pep456

### Slide 8
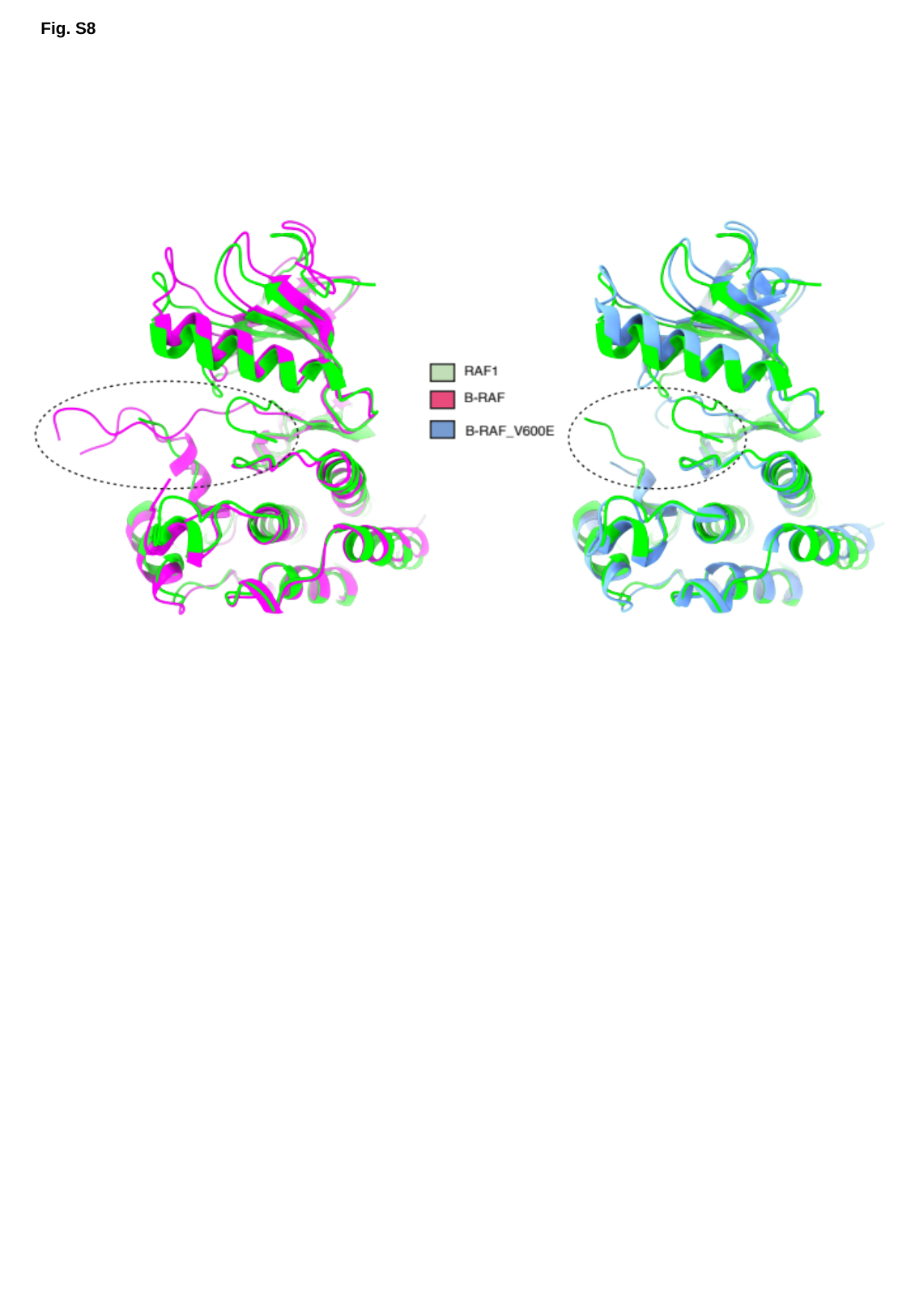

Fig. S8
